# The interaction landscape between transcription factors and the nucleosome

**DOI:** 10.1101/240598

**Authors:** Fangjie Zhu, Lucas Farnung, Eevi Kaasinen, Biswajyoti Sahu, Yimeng Yin, Bei Wei, Svetlana Dodonova, Patrick Cramer, Jussi Taipale

## Abstract

Nucleosomes cover most of the genome and are thought to be displaced by transcription factors (TFs) in regions that direct gene expression. However, the modes of interaction between TFs and nucleosomal DNA remain largely unknown. Here, we use nucleosome consecutive affinity-purification systematic evolution of ligands by exponential enrichment (NCAP-SELEX) to systematically explore interactions between the nucleosome and 220 TFs representing diverse structural families. Consistently with earlier observations, we find that the vast majority of TFs have less access to nucleosomal DNA than to free DNA. The motifs recovered from TFs bound to nucleosomal and free DNA are generally similar; however, steric hindrance and scaffolding by the nucleosome result in specific positioning and orientation of the motifs. Many TFs preferentially bind close to the end of nucleosomal DNA, or to periodic positions at its solvent-exposed side. TFs often also bind nucleosomal DNA in a particular orientation, because the nucleosome breaks the local rotational symmetry of DNA. Some TFs also specifically interact with DNA located at the dyad position where only one DNA gyre is wound, whereas other TFs prefer sites spanning two DNA gyres and bind specifically to each of them. Our work reveals striking differences in TF binding to free and nucleosomal DNA, and uncovers a rich interaction landscape between the TFs and the nucleosome.

---

Simple prokaryotic organisms such as *E.coli* have relatively small genomes, which are often organized into a circular chromosome consisting of a single DNA molecule. Their genes are regulated by TFs that directly bind to the free DNA molecule and influence transcriptional activity. Eukaryotic genomes, however, are much larger, and need to be packaged more efficiently inside the nucleus. The packaging is accomplished by a specific class of basic proteins, the histones, which exist as an octameric complex and bind to the DNA backbone, forming nucleosomes^1-4^. In a canonical nucleosome, a 147 bp segment of DNA is wrapped around the histone octamer in a left-handed, superhelical arrangement for a total of 1.65 turns, with the DNA helix entering and exiting the nucleosome from the same side of the histone octamer. The two DNA gyres are paralleling each other except at the position located between the entering and the exiting DNA, where a dyad region of ~15 bp contains only a single DNA gyre. The nucleosome is 2-fold pseudo-symmetric with respect to a dyad axis at the center of the dyad region. Approximately 70% of eukaryotic DNA is packaged into nucleosomes, separated from each other by free DNA linker sequences of 10–80 bp^5-7^.

The nucleosome presents a significant barrier for binding of other proteins such as RNA polymerases to DNA^8-14^. As a consequence, the presence of nucleosomes can have a negative effect on gene expression. Similarly, most TFs are thought to be unable to bind to nucleosomal DNA, and TF binding sites in the genome are usually depleted of nucleosomes^15-17^. However, it is thought that a specific class of TFs, the pioneer factors, can access nucleosomal DNA, and assist the binding of other TFs to nearby sites^18-22^. Many TFs that have essential roles in development and cell reprogramming are pioneer factors^23,24^. Two different mechanisms have been suggested to be responsible for the pioneering activity: mimicking the linker histones^25^ and/or targeting a partial TF motif that is accessible on nucleosomal DNA^26^.

Nucleosomes can also indirectly induce cooperativity between multiple TF binding events^27-31^. This cooperation can occur in the absence of direct TF-TF interactions^32^, allowing multiple weak binding events to dissociate nucleosomes, resulting in a preferred range of spacings between the two TF binding sites^33^. Consistently, in higher eukaryotes, most occupied TF binding sites are clustered to short genomic regions^34-37^.

Despite the importance of the nucleosome in both chromatin organization and transcriptional control, the effect of nucleosomes on TF binding has not been systematically characterized. This is in part because the sites bound by both a TF and a nucleosome are difficult to identify in cells, as the methods to map cellular TF binding locations are imprecise. Furthermore, TF-nucleosome complexes that activate chromatin remodeling in cells are expected to be unstable, and thus hard to capture experimentally. In our recent work, we found that scaffolding by DNA results in a large number of interactions between transcription factors^38^. Given that the DNA scaffold is bent and partially blocked by the nucleosome, it is likely that nucleosome occupancy will also have a major effect on TF-DNA binding.

## RESULTS

### Nucleosome CAP-SELEX

To determine the effect of nucleosome on TF-DNA binding, we adapted our previous Consecutive Affinity-Purification SELEX (CAP-SELEX) method^38^ to include nucleosome reconstitution. We name this approach Nucleosome CAP-SELEX (NCAP-SELEX). We designed two types of SELEX ligands, a 147 bp (lig147) and a 200 bp (lig200) ligand, containing 101 bp and 154 bp randomized regions, respectively. In the NCAP-SELEX assay (**Fig. 1a**), recombinant histone octamers containing tagged H2A proteins (**Extended Data Fig. 1a, b**) are first loaded onto the DNA ligands (**Extended Data Fig. 1c, d**) in 384-well microplates by decreasing the salt concentration in a stepwise fashion (see **Methods** and Dyer *et al*.^39^). Subsequently, the nucleosomes are purified using magnetic beads. Eluted nucleosomes are incubated with TFs having an orthogonal tag, and the TF-bound species are subsequently pulled-down. The bound DNA is then amplified using PCR, and the entire process is repeated for a total of five times. To determine whether the TF binding induces dissociation of nucleosomes, the nucleosomes are recaptured after the final cycle (**Fig. 1a**). Both the nucleosome-bound and unbound DNA of the final cycle, as well as input DNA and DNA from the earlier cycles are then sequenced using a massively parallel sequencer.

**Figure 1.**
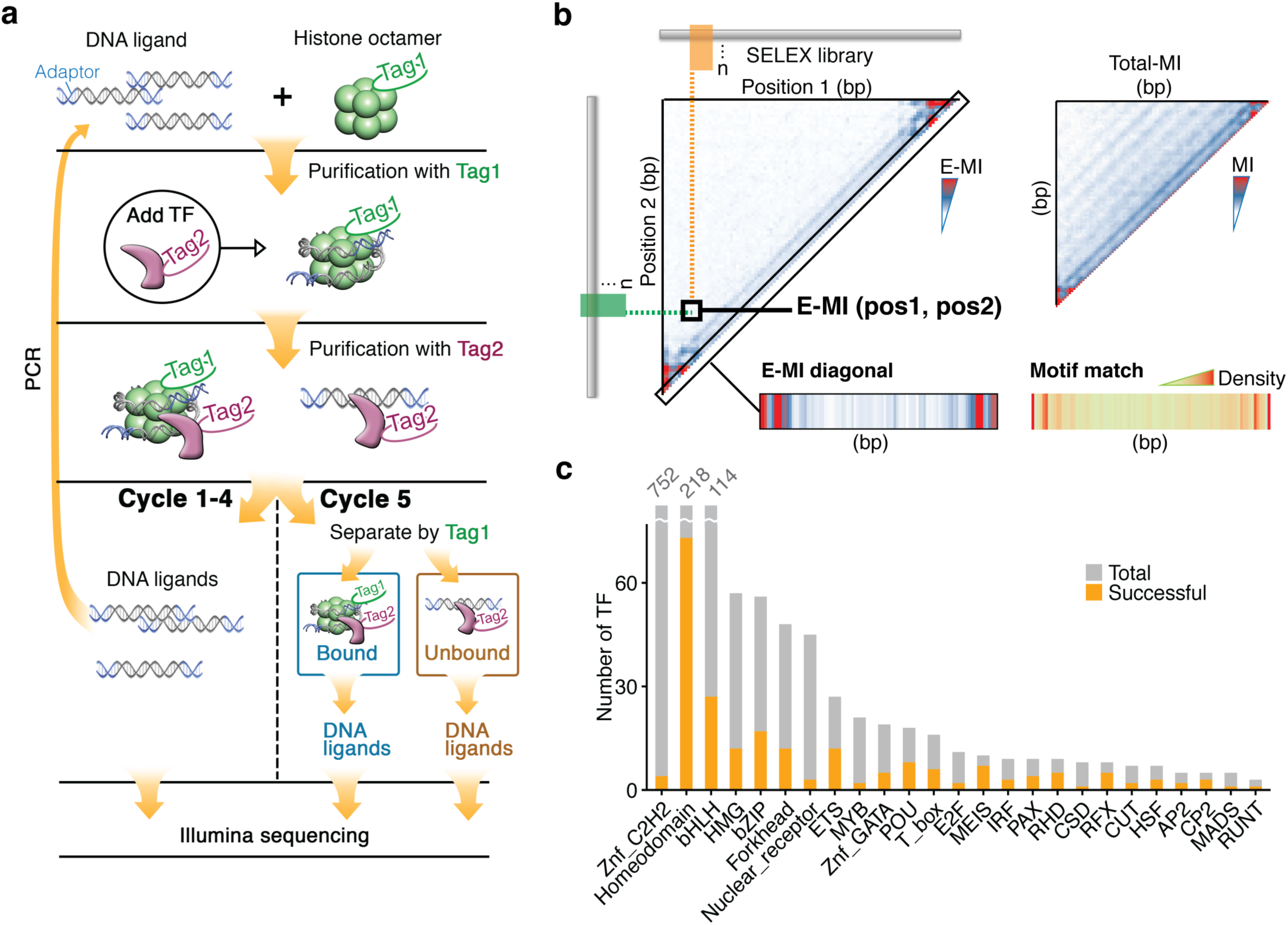
Nucleosome CAP-SELEX. **a**, Schematic representation of NCAP-SELEX. The DNA ligands for SELEX contain a randomized region (grey) with fixed adaptors (blue) at both ends. The protocol first selects for ligands that are favored by the nucleosome, and then from the nucleosome-bound ligand pool selects for ligands that bind to a given TF. The orthogonal tagging of histone H2A (tag1) and TFs (tag2) enables the consecutive affinity purification. In the last (5th) cycle, the TF-bound DNA ligands are further separated into nucleosome-bound and unbound libraries, **b**, TF-signal analysis by E-MI. For the library enriched by each TF, E-MI (Mutual Information between the most Enriched 3-mer pairs) is calculated pairwise between all non-overlapping 3-mer columns (left triangle). When analysing TF signals, we chose E-MI instead of total-MI (right triangle) because total-MI detects also signals from the nucleosome (the stripes with 10-bp intervals). The diagonal of the E-MI plot (bottom left) is most informative, and is generally in line with the motif-matching result (bottom right), **c**, Family-wise coverage of successful TFs.

To determine the effect of nucleosomes on TF binding, the ligand sequences were analyzed computationally using motif matching, *de novo* motif discovery and mutual information pipelines as illustrated in **Extended Data Fig. 1e**. In most analyses, we estimate TF signals using an approach that is based on the mutual-information (MI) between 3-mer distributions at two non-overlapping positions of the ligand (**Fig. 1b**). The underlying rationale is that if a binding event contacts two positions of a SELEX ligand at the same time, the 3-mer distributions at these two positions would be correlated in the enriched library, with the joint distribution favoring the 3-mer combinations that form the high-affinity sites. This biased joint distribution would then be detected as an increase in MI between the positions. Such an approach has multiple advantages: it operates without previous knowledge of TF specificities, enables facile comparison of selectivity between different TFs, and pinpoints the positions where DNA interacts with the TFs.

For the library enriched by each TF, we calculated MI between all pairwise combinations of positions, and represented the results as a 2D heatmap (**Fig. 1b**). In the heatmap showing MI from all 3-mer pairs (total MI; **Fig. 1b**, right), stripes with ~ 10 bp spacing are visible in addition to the TF signals. These stripes reflect the nucleosome signal, as histones contact DNA at ~ 10 bp intervals^40-43^. To focus more on the TF signals, we further developed a measure that considers only the MI between the ten most enriched non-overlapping 3-mer pairs (E-MI; **Fig. 1b**, left). As TFs rarely bind or cooperate across a large span of DNA, their signals are usually located at the diagonal of the 2D E-MI map. Therefore, the E-MI diagonal (**Fig. 1b**, bottom left) captures most of the footprints of TFs on the ligands, and corresponds well with the distribution of matches to the TFs’ motifs (**Fig. 1b**, bottom right).

We performed NCAP-SELEX using 413 human TF extended DNA binding domains (eDBDs) (details are given in **Supplementary Table 1** and **Methods**). The selected TFs covered 28% of the high-confidence TFs reported by Vaquerizas *et* al.^44^. Among the tested TFs, 220 eDBDs were successful (**Fig. 1c;** see **Methods** for details).

### Nucleosome inhibits TF binding

We next analyzed the effect of nucleosome on TF binding by clustering the E-MI diagonal signals from the lig200. The result reveals that the binding of almost all TFs to DNA is inhibited or spatially restricted by the presence of a nucleosome (**Fig. 2a**). The lig200 can accommodate only one nucleosome, but allows multiple positions for it (**Fig. 2a**, schematic at the center). The nucleosome occupancy is thus expected to increase towards the center of the lig200. The penetration of the E-MI diagonal signal into the center, in turn, reflects the ability of each TF to bind to nucleosomal DNA. Analysis of this data revealed that TFs from the same family tend to cluster together based on the E-MI diagonal (**Fig. 2a**; SOX TFs indicated as an example). However, the extent of the penetration varied strongly between the TFs (**Fig. 2b**). For example, SREBF1 and 2, and RFX3 (**Fig. 2c**, left) only show E-MI signal at the extreme ends of the ligand, suggesting that they have weak affinity towards nucleosomal DNA relative to free DNA. In contrast, TFs such as VSX1, ARX, and SOX12 display stronger signal near the center (**Fig. 2c**, right), and are thus more capable of binding to the nucleosome-occupied regions.

**Figure 2.**
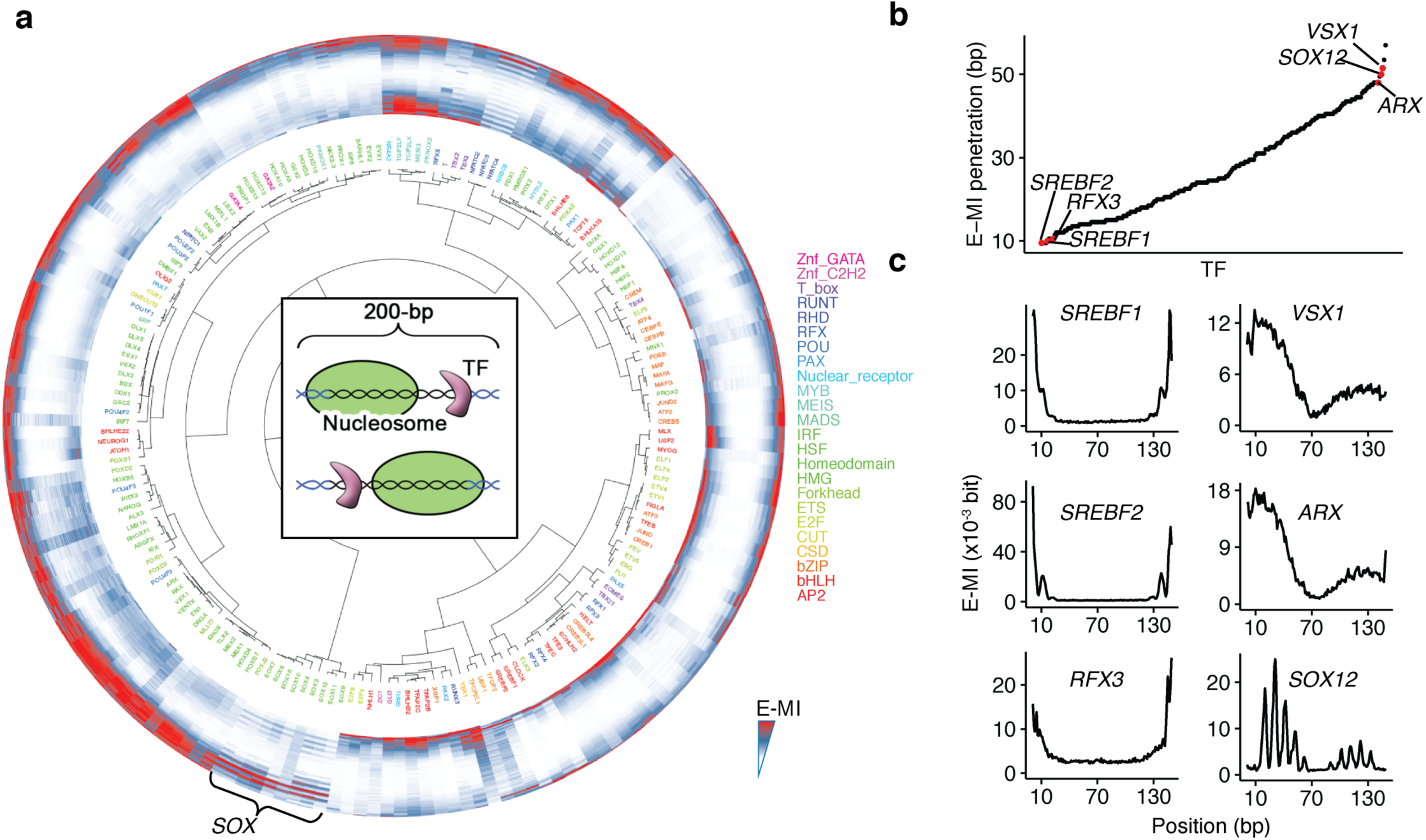
Nucleosomal DNA is less accessible for TFs than free DNA. **a**, Hierarchical clustering of the E-MI diagonals for NCAP-SELEX with the 200-bp ligand (lig200). The E-MI diagonal for each TF is oriented radially. The names of the TFs are colored by family with the coloring scheme indicated on the right. TFs from the same family tend to be clustered together (e.g., SOX, indicated). The illustration at the center of the dendrogram schematically represents TF’s binding on lig200. Note that almost all TFs have lower E-MI towards the center of lig200, indicating their lower affinity to nucleosomal DNA compared with free DNA. The E-MI diagonals are scaled for each TF. **b**, E-MI penetration of individual TFs on lig200. TFs are ordered according to their E-MI penetration depth towards the center of the ligand. This order reflects TFs’ ability to bind nucleosome-occupied DNA. Six TFs representing either of the two extremes are colored red and exemplified in **(c). c**, The diagonal of E-MI for TFs with highest/lowest E-MI penetrations. Left: TFs with lowest E-MI penetrations; right: TFs with highest E-MI penetrations.

### TFs can bind sequences located on both nucleosomal DNA gyres

For lig200 libraries, we next analyzed the entire 2D E-MI signals for individual TFs. This analysis resulted in identification of a specific binding mode for T-box TFs on nucleosomal DNA. Binding of brachyury (T) to nucleosomal DNA resulted in two prominent E-MI signals (**Fig. 3a**, the heatmap). One was located at the E-MI diagonal, i.e. observed between adjacent 3-mers, whereas the other resulted from 3-mers located ~ 80 bp from each other. The first signal is due to the binding of T to nucleosomal DNA using motifs similar to those found on free DNA (**Fig. 3a**, Mode 1). The second is associated with an approximately 80-bp-long motif (**Fig. 3a**, Mode 2), indicating a dimeric binding spanning the two gyres of the nucleosomal DNA (**Fig. 3b**). We next compared the bound and unbound libraries of the last cycle, and found that the signal for Mode 2 is stronger on the ligands that remained bound to the nucleosome (**Fig. 3c**), indicating that the gyre-spanning binding stabilizes mononucleosomes against dissociation. The Mode 2 binding is also observed for another T-box factor, TBX2, and is not detected on free DNA (**Extended Data Fig. 2a, b**).

**Figure 3.**
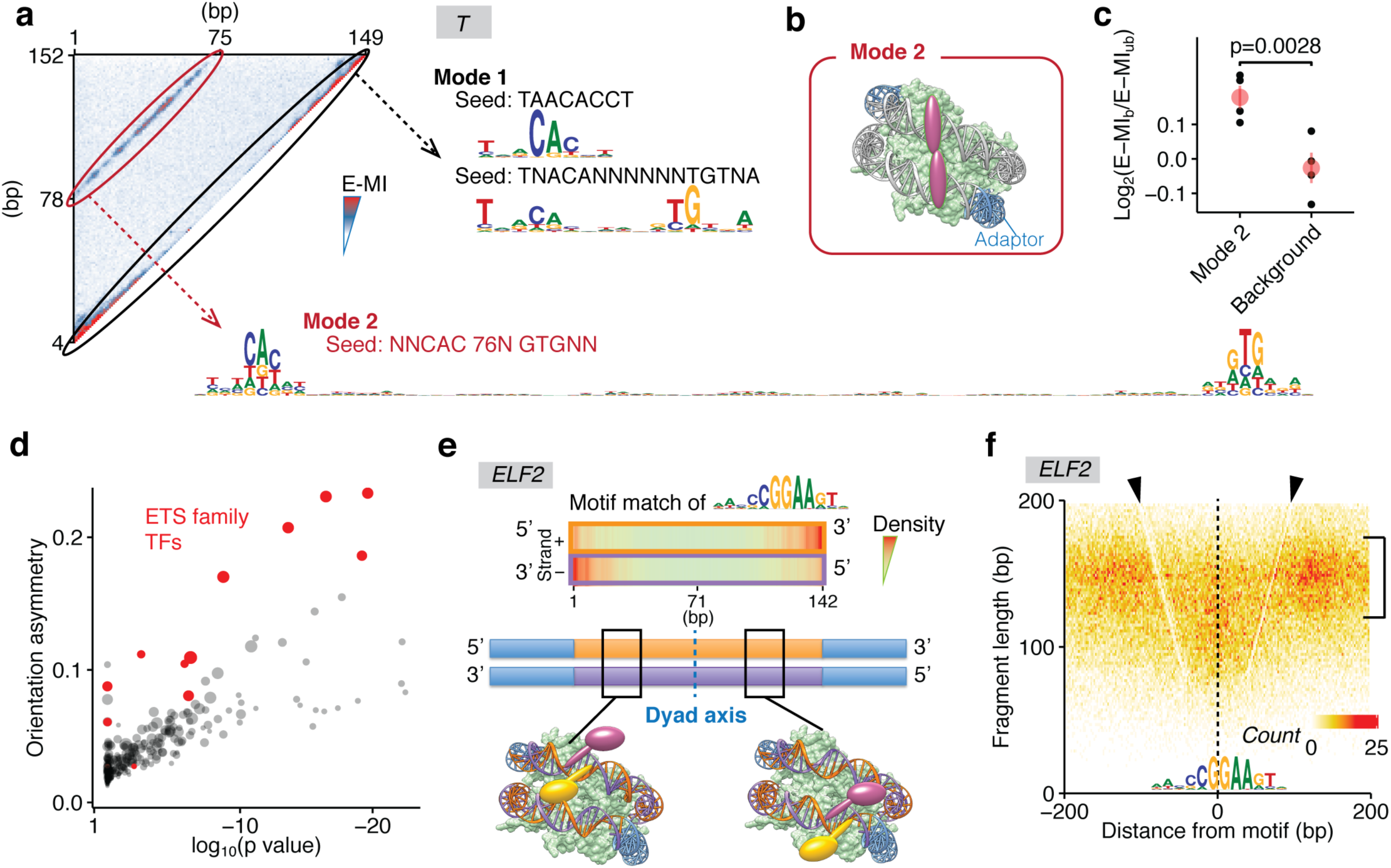
Nucleosome allows binding that spans two gyres and breaks the rotational symmetry of DNA. **a**, Two modes used by T (brachyury) to bind nucleosomal DNA. The heatmap shows the pairwise E-MI for all combinations of positions on the 200-bp ligand. The Mode 1 signal near the diagonal gives motifs similar to those seen on the free DNA. The Mode 2 signal corresponds to a ~80-bp-long motif. Mode 1 is inhibited by higher nucleosome occupancy towards the center whereas Mode 2 gets stronger in the middle. Seed for each motif is also indicated, **b**, Schematic representation of TFs that bind the two gyres of nucleosomal DNA at the same time. **c**, Mode 2 binding stabilizes nucleosome from dissociation. Log2 ratio of E-MI between the bound and unbound libraries (cycle five, four replicates) of T is calculated for both the Mode 2 binding and the background E-MI level (see **Method** for details). The bound library has stronger Mode 2 binding but similar background. Each point indicates a replicate. Data are mean ± s.d.; two-sided t-test was used, 95% CI, 0.097–0.202. **d**, Orientational asymmetry of individual TFs. For each TF, the asymmetry is evaluated by the binding energy difference between the two relative orientations, averaged for 40 non-palindromic 8-mers that are most enriched in the TF’s NCAP-SELEX library; significance of the asymmetry is also tested to obtain the p value (see **Method** for details). Most of the ETS-family TFs (red) show a prominent orientational asymmetry. Dot size represents the extent of signal enrichment in each TF’s NCAP-SELEX library, **e**, The orientational asymmetry of ELF2. The ETS factor ELF2 has different motif density distributions for the two strands of nucleosomal DNA (top panel). This is because at a specific position, TFs (magenta and yellow, bottom panel) that respectively bind to motifs on different DNA strands (purple and orange, blue for constant adaptor region) will differ in their surrounding chemical environments. For motifs that locate on different DNA strands and equidistant from the dyad, their chemical environment for binding will be identical due to the rotational symmetry of the nucleosome (with respect to the dyad axis), e.g., the magenta TF in the left model has the same chemical environment as the yellow TF in the right model. As a result, the motif densities on two DNA strands are different but symmetric to each other with regard to the dyad. **f**, The asymmetric MNase fragment profile around genomic ELF2 sites. ELF2 motif matches within ChIP-seq peaks were positioned at the center. Nearby MNase fragment counts are summarized with 2 × 2 bins according to their lengths and center positions. Nucleosome distribution near the ELF2 sites are reflected by the signal intensity of the ~150 bp fragments (bracket). The V-shaped lines with a lower signal intensity (arrowheads) reveal the footprint of the TF, which is asymmetric.

Interestingly, the scaffolding effect of the nucleosome also leads to TF binding modes that contact nucleosomal DNA at positions spaced by approximately 40 bp (e.g. TBX2 and ETV, as shown in **Extended Data Fig. 2b, c**). This effect is position-specific, with one binding event being observed near the dyad, and the other(s) on the opposite side of the nucleosome, with the two contacts separated by ~180°. As the individual TFs are located far from each other in this binding mode, the binding pattern suggests that the nucleosome may have two allosteric states or may form a higher order complex with these TFs.

Taken together, these results reveal that some TFs can interact with both DNA gyres on the nucleosome, and suggest that nucleosome can generate novel composite TF-binding sites on DNA by promoting spatial proximity of DNA sites that are located more distally on free DNA.

### Nucleosome context breaks the rotational symmetry of DNA

As DNA is double-stranded, TFs can bind to it in two different orientations. For TFs that bind non-palindromic sites, their binding orientation can be determined from the bound sequences. In analysis of motif matches on lig200, we noted that some TFs’ motifs displayed a bias of matches in one orientation at the 5’ end, and in the other orientation at the 3’ end of the ligand. That is, these TFs have a preferred orientation relative to the nucleosome. We systematically examined this asymmetric effect between binding orientations by comparing the strand-wise distributions of top 8-mers (**Fig. 3d**, **Extended Data Fig. 3a**, see also **Methods** for details).

Both the extent of the orientational asymmetry and the associated p-value (**Fig. 3d**) revealed that many ETS factors displayed strong orientational preferences. ELF2 is shown in **Fig. 3e** as an example; its motif distributions (**Fig. 3e**, upper panel) and top 8-mer distributions (**Extended Data Fig. 3b**) display strong orientational preference. CREB factors also show considerable orientational preference towards the nucleosome in NCAP-SELEX (**Extended Data Fig. 3c**). The orientational asymmetry induced by the nucleosome can be explained by the fact that DNA is rotationally pseudosymmetric, and this symmetry is broken by the presence of the nucleosome (**Extended Data Fig. 3d**), leading to a different local environment for a TF bound at the same position of DNA in opposite orientations (**Fig. 3e**, red and yellow ovals). Depending on its orientation, a particular side of a TF will be in proximity with either the second gyre of nucleosomal DNA, or the histone proteins.

The distributions of motif matches in the two strands were symmetric with regard to the dyad position of the nucleosome. This, in turn, is a consequence of the pseudo 2-fold symmetry of the nucleosome; two binding sites in different orientations will share an identical configuration when they locate at opposite sides of the dyad, and have an equal distance to the dyad (**Fig. 3e**, models in the lower panel).

To determine whether the directional binding of TFs to a nucleosome is also observed *in vivo*, we performed MNase digestion followed by paired-end sequencing for the human colorectal cancer cell line LoVo. We then visualized the distribution of MNase fragments around directional ELF2 motif matches within ELF2 ChIP-seq peaks from Yan et al.^37^ (**Fig. 3f**). As described previously^45^, this visualization reveals nucleosomes near the TF sites due to enrichment of fragments whose size corresponds to a single nucleosome. The footprint of the TF is also seen as a V-shaped line having lower signal intensity (arrowheads in **Fig. 3f**). This analysis shows that both the nucleosome distribution and the TF footprint size are asymmetric with respect to the ELF2 sites. For the specified motif direction, the footprint of ELF2 is more distinct downstream of the nucleosome than upstream of it. This implies a more stable binding of ELF2 downstream of the nucleosome, which is in accordance with the motif match analysis from the ELF2 NCAP-SELEX data (**Fig. 3e**). The MNase analysis also indicated that nucleosome occupancy is lower upstream than that downstream of ELF2 sites. This pattern suggests that the more stable binding of ELF2 downstream of the nucleosome displaces the nucleosome or pushes it upstream. Similar to ELF2, the binding profile of ELF1 is also asymmetric with regard to nucleosome both in SELEX and *in vivo* (**Extended Data Fig. 3e**).

### Nucleosome induces positional preference to TF binding

We next analyzed the positional preference of TF binding on nucleosomal DNA using the short lig147 ligand. Because its 147-bp length exactly matches the preferred length of nucleosomal DNA, the nucleosome is expected to be uniquely positioned at the center of lig147. Therefore, the relative positioning of the TFs with respect to the nucleosome can be inferred at a higher resolution than using lig200. To determine the positional preference, we first checked whether TFs’ motifs on nucleosomal DNA are different from their motifs on free DNA. For this purpose, we compared the most enriched 9-mer sequences for each TF, between its lig147 libraries enriched either in the presence and absence of the nucleosome (**Extended Data Fig. 4a)**. The result shows that most TFs bind to similar 9-mers under both conditions, suggesting that TFs are binding nucleosomal DNA without significant specificity changes. However, consistent with earlier observations^26^, we also found few cases where the binding specificities of the TFs were detectably different on nucleosomal DNA (**Extended Data Fig. 4b**).

Analysis of TF binding to lig147 revealed several types of positional preference (**Fig. 4a**), which we classified into three major classes (**Fig. 4a**): (1) End binders; these TFs tend to prefer positions towards the end of the ligand. All tested bZIP factors belong to this class (**Fig. 4b**), e.g., CEBPB (**Fig. 4c, Extended Data Fig. 5a**). This preference might be explained by the “breathing”, i.e. the spontaneous partial detachment of nucleosomal DNA^1,46,47^, which occurs more frequently towards the entry and exit of nucleosomal DNA (**Fig. 4d**). (2) Periodic binders; these TFs tend to bind periodic positions on nucleosomal DNA. This periodicity is likely induced by the contacts of histones to DNA at 10-bp intervals. (3) Dyad binders; these TFs prefer to bind nucleosomal DNA near the dyad position. In addition to these three classes, we also identified a “mixed” class (**Fig. 4a**) where TFs show E-MI diagonal characteristics of both the end binder and the periodic binder class. TFs behaved consistently for lig147 and lig200 according to the binder classification (**Extended Data Fig. 5b**). Compared to the end binders, the periodic binders and dyad binders displayed deeper penetration of E-MI signals into the center of the ligands (**Extended Data Fig. 5b**); they are thus more capable to bind nucleosomal DNA.

**Figure 4.**
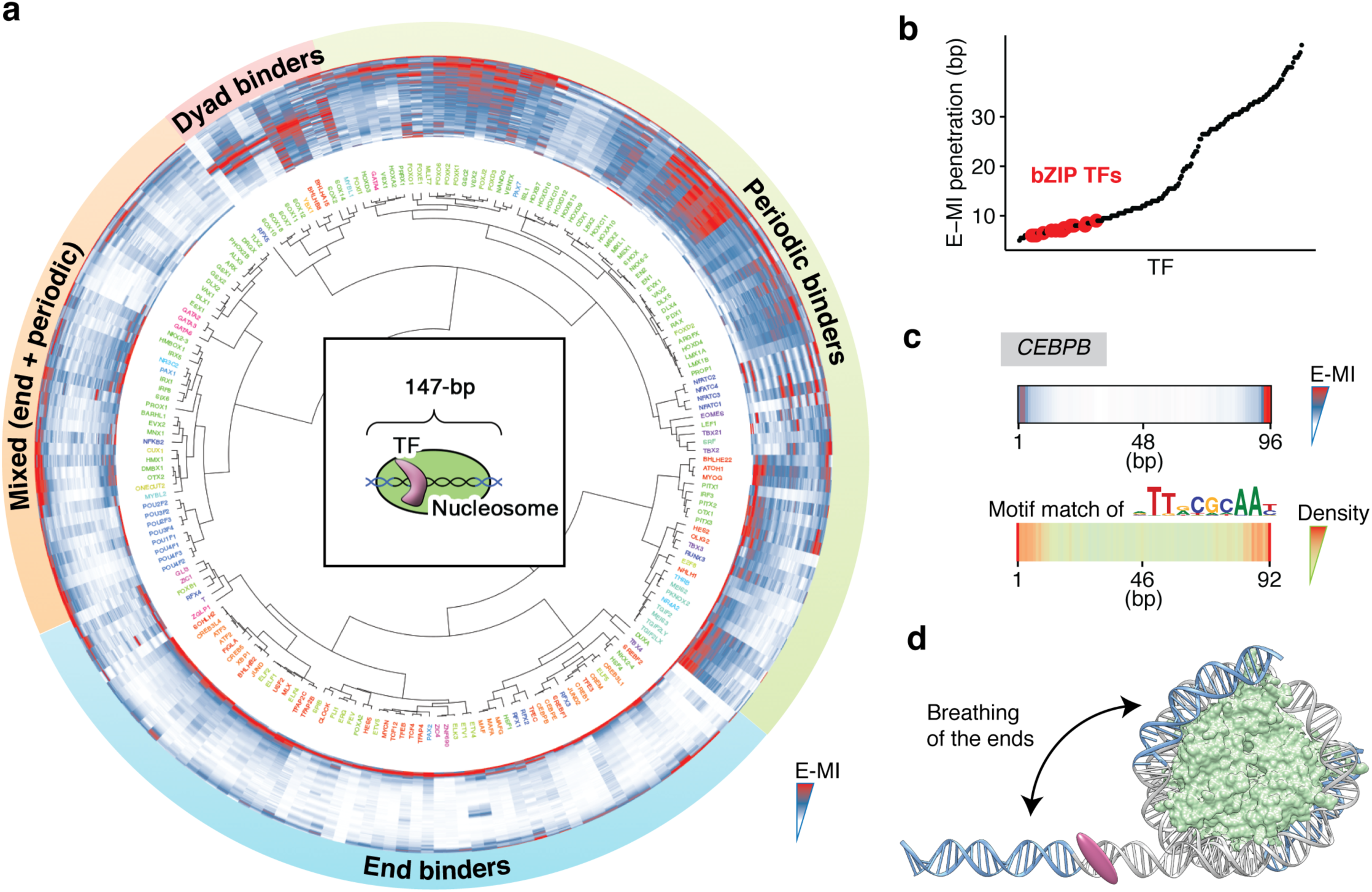
Nucleosome induces positional preference to TF binding. **a**, Hierarchical clustering of the E-MI diagonals for NCAP-SELEX with the 147-bp ligand (lig147). The coloring scheme is the same as that in **Fig. 2a**. In the center of the dendrogram, the schematic shows that nucleosome is positioned uniquely on lig147. TFs are assigned to three separate classes and a mixed class. E-MI diagonal is scaled for each TF. **b**, E-MI penetration of each TF on lig147. All examined bZIP TFs are marked with red. Their low penetrations indicate an end preference. **c**, E-MI diagonal and motif matching results for the bZIP factor CEBPB. **d**, Schematic representation showing a TF is preferring the ends of nucleosomal DNA due to breathing. Both the two ends of nucleosomal DNA will breath but only one is illustrated here for clarity.

### Binding at the outward-facing side of the DNA helix

Half of the circumference of nucleosomal DNA is in close proximity of the histones. As DNA is helical, equivalent positions that could be accessible to TFs are thus located at ~10 bp intervals. Accordingly, we found that many TFs prefer to bind to positions located ~10 bp apart on nucleosomal DNA (**Fig. 4a**, periodic binders). We studied this effect using the lig147 libraries. By applying a Fast Fourier Transform (FFT) to the E-MI diagonals, we obtained the strength and phase of the ~10 bp periodicity for the TFs (**Fig. 5a**). The result shows that the overall periodicity of E-MI is stronger for the NCAP-SELEX library compared to the free-DNA HT-SELEX library (**Fig. 5a**, bottom). Due to the binding specificity of nucleosome, an increased periodicity was also observed with the counts of dinucleotides (e.g. TA) along the ligand (**Extended Data Fig. 6a**). TA-enriched positions on nucleosomal DNA correspond to positions where histones contact DNA^4,42^, which are also positions where the DNA major groove is facing towards the solvent. The periodicity of TA for all experiments had a similar phase (**Extended Data Fig. 6a**), suggesting that in NCAP-SELEX, the nucleosomes reconstituted for all TFs shared a similar rotational position on the DNA ligand. In contrast, the phase of the E-MI periodicity is much more dispersed (**Fig. 5a**). This dispersion is consistent with the preference of TFs towards the minor and major grooves of DNA (**Fig. 5b, c**).

**Figure 5.**
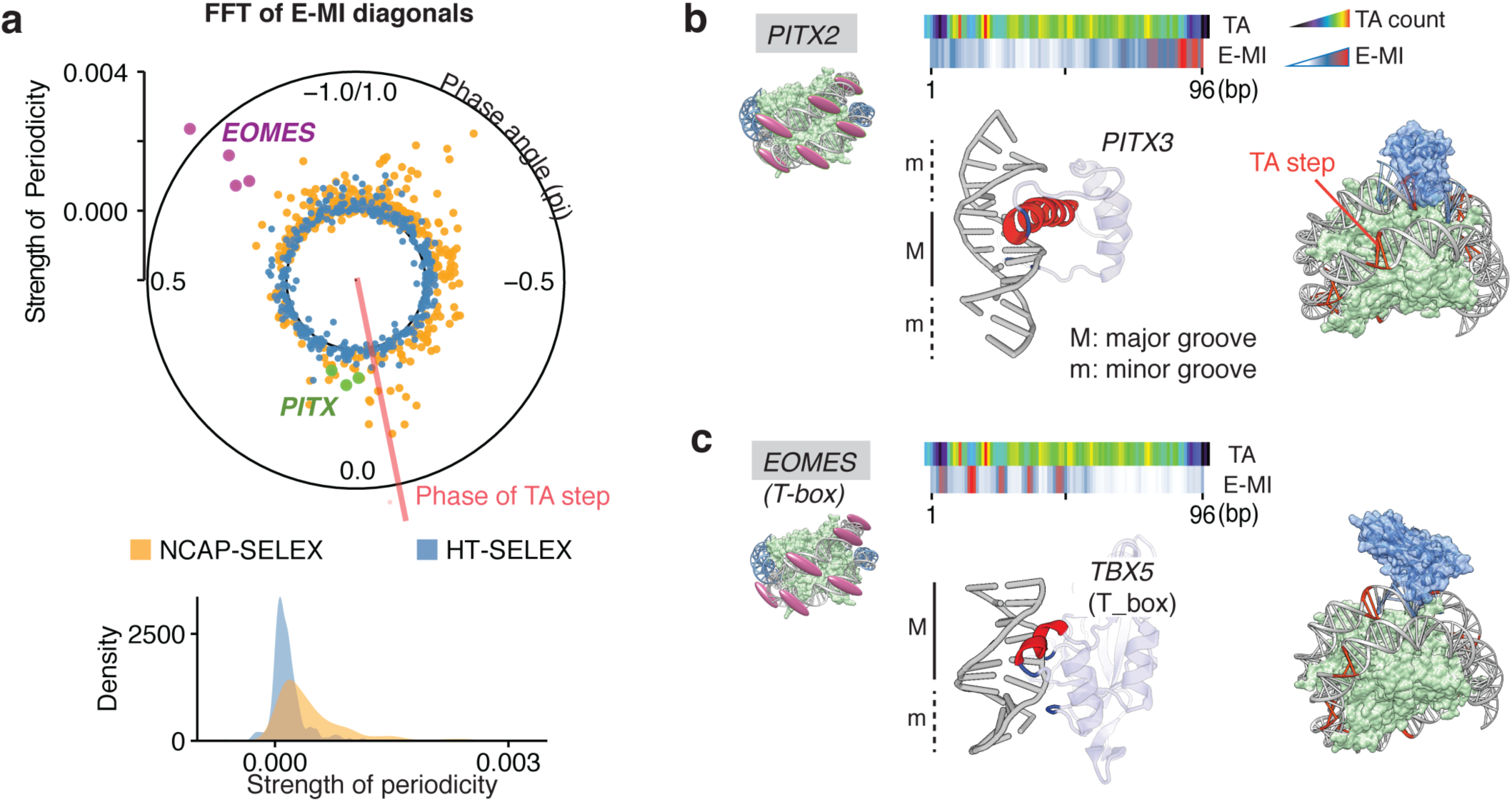
Periodic binding of TFs to the major or minor grooves facing outwards. **a**, Strength and phase of the ~10 bp periodicity for individual TFs. The polar plot shows the strength and phase of the periodicity derived from the FFT of E-MI diagonals; data for both the NCAP-SELEX (orange; nucleosomal DNA) and the HT-SELEX (blue; free DNA) are shown. Each dot represents one SELEX library. EOMES (magenta, four replicates) and PITX (green for PITX1, 2, 3) have opposite phases; they are exemplified in **(b, c)**. The phase of the TA dinucleotide (red line, median value of all NCAP-SELEX libraries) is also shown to indicate where histones contact nucleosomal DNA^42^. Bottom: density plot of the periodicity strength for all TFs. **b**, Major groove binder prefers exposed major grooves on nucleosomal DNA. The E-MI diagonal of PITX is in phase with the TA peaks along the ligand. The structure of PITX (PDB 2lkx, visualized with DNAproDB^83^) also show contacts with DNA principally in the major groove (M). The base-contacting helices (red) and loops (blue) are indicated. Cartoon representation to the right shows that the steric hindrance is minimal when PITX (blue) binds in phase with TA (orange) on the nucleosome structure (PDB 3ut9). **c**, Minor groove binder prefers exposed minor grooves (m) on nucleosomal DNA. The E-MI diagonal of EOMES (T-box) is out of phase with the TA peaks, suggesting it binds positions where nucleosomal DNA’s minor groove is facing outside. The TBX5 (T-box) structure (PDB 2×6v) also shows contacts with DNA principally in the minor groove. Cartoon representation to the right shows that the steric hindrance is minimal when TBX5 (blue) binds out of phase with TA (orange) on the nucleosome structure (PDB 3ut9).

For example, PITX and EOMES prefer almost opposite phases of nucleosomal DNA (**Fig. 5a**), respectively in phase and out of phase with the TA dinucleotide (**Fig. 5b, c**; the heatmaps). Consistently, the structural analysis indicated their different groove preference: PITX contacts DNA principally by insertions into the major groove (structure in **Fig. 5b**)^48^, whereas the T-box TFs principally contact DNA via the minor groove (structure in **Fig. 5c**; see also the references^49,50^). Because the E-MI measure detects the most enriched 3-mer pairs, high E-MI signal usually occurs at positions that correspond to direct TF amino-acid to DNA contacts. Thus, TFs that bind to the major groove tend to show E-MI maximums in phase with TA, and TFs that bind to the minor groove commonly display E-MI maximums out of phase with TA, as seen in **Fig. 5b** and **5c**. Such patterns of TF binding minimize the steric conflict between TF and the histones (cartoon of TF-nucleosome complex in **Fig. 5b, c** and **Extended Data Fig. 6b**).

The periodic pattern of E-MI diagonal agrees with the motif matching result (**Extended Data Fig. 6c**). The periodic availability of DNA for TF binding also imposes a ~10 bp periodicity on dimer spacing patterns (**Extended Data Fig. 6d**) for individual TFs that can bind to the outward-facing DNA. However, in most cases such binding appears not to be cooperative, based on the fact that the observed frequency of ligands with two motifs can be well estimated by the frequency of ligands that contain only one motif (data not shown). Taken together, our results indicate TFs tend to bind to the outward-facing side of nucleosomal DNA, as expected from steric considerations.

### Binding near the nucleosomal dyad

Analysis of the positional preference of TFs on nucleosomal DNA also revealed that the region around the nucleosomal dyad is strongly preferred by a few TFs. For example, RFX5 shows the strongest binding around the dyad position of lig147 (**Fig. 6a**, **Extended Data Fig. 7a**). Also, multiple SOX TFs show a preference for binding to DNA near the dyad (**Fig. 6b**). Distinct from other regions of nucleosomal DNA, the dyad region contains only a single DNA gyre (**Fig. 6c**), and the histone disk is thinnest there^41,51^. These features of the dyad DNA reduce the steric barrier for TF binding, and could allow TFs that bend DNA upon binding (such as SOX proteins^52^) to deform DNA relatively easily.

**Figure 6.**
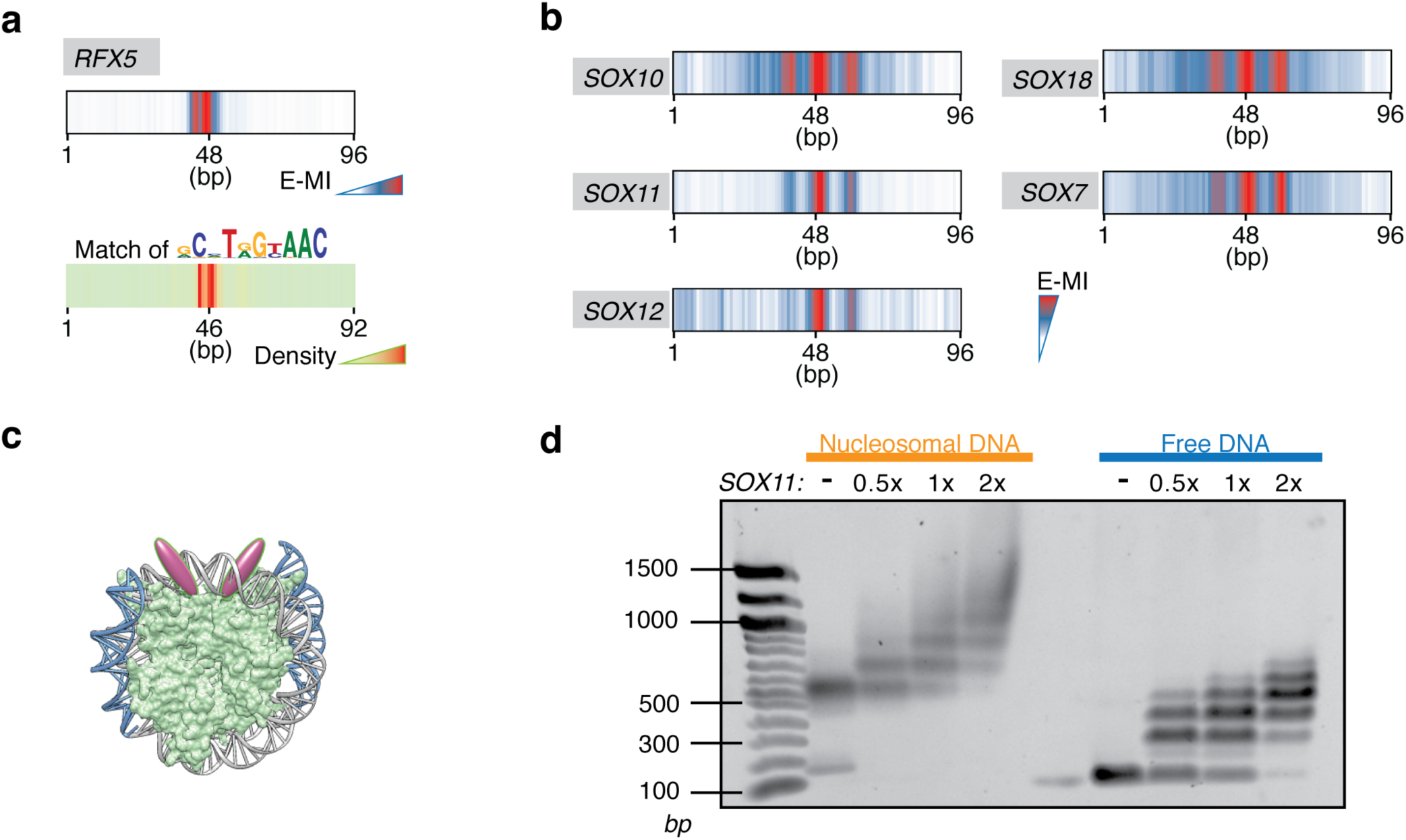
Binding near the dyad axis. **a**, E-MI diagonal and motif matching results for RFX5. **b**, E-MI diagonal of SOX family TFs showing their preferred binding around the dyad. **c**, Schematic representation of TFs that prefer to bind around the dyad. **d**, EMSA of SOX11 complexes with nucleosome and with free DNA. Nucleosome is reconstituted and purified using a modified Widom 601 sequence, which contains a SOX11 binding sequence (extracted from cycle 4 SELEX library) embedded close to the dyad. Each 40 μl reaction contains 1 μg DNA, together with SOX11 protein at a molar ratio of 0, 0.5, 1, 2 (indicated on top of each lane) to DNA.

Binding of SOX11 to sites near the dyad of nucleosome was validated with Electrophoretic Mobility Shift Assay (EMSA). Nucleosomes containing a SOX11 binding sequence identified in the NCAP-SELEX experiment were incubated with increasing amounts of purified SOX11 eDBD. The clear super-shift confirmed the binding of SOX11 to the nucleosome (**Fig. 6d**). The result also indicates that SOX11 does not dissociate the nucleosome upon binding.

### TFs and their binding positions differ in the ability to dissociate the nucleosome

To determine whether TF binding affects the stability of the nucleosome, we performed an additional affinity capture step to separate the nucleosome-bound and dissociated DNA (unbound) after the last NCAP-SELEX cycle (**Fig. 1a;** lig147). As a control experiment, we also allowed the last-cycle nucleosome to dissociate without the presence of TFs. TFs whose binding leads to nucleosome dissociation are expected to have more and stronger binding sites in the unbound library compared to the bound library. Conversely, TFs that stabilize the nucleosome will show the reverse. To evaluate each TF’s effect on the stability of the nucleosome, the differential E-MI between its bound and unbound libraries was calculated. Control experiments lacking TFs showed very little effect (**Extended Data Fig. 8a**), whereas in the presence of TFs, clear differences in E-MI signals were observed (**Fig. 7a** and **Extended Data Fig. 8a**). We found that most TFs (e.g. CDX1) have stronger E-MI in the unbound library compared to that of the bound library (**Fig. 7a, b**), suggesting that they can facilitate nucleosome dissociation upon binding. However, we also identified a few exceptional TFs whose binding stabilized the nucleosome. These include the T-box TFs, such as TBX2. All three TBX2 replicates had higher E-MI in the bound library (**Fig. 7b**).

**Figure 7.**
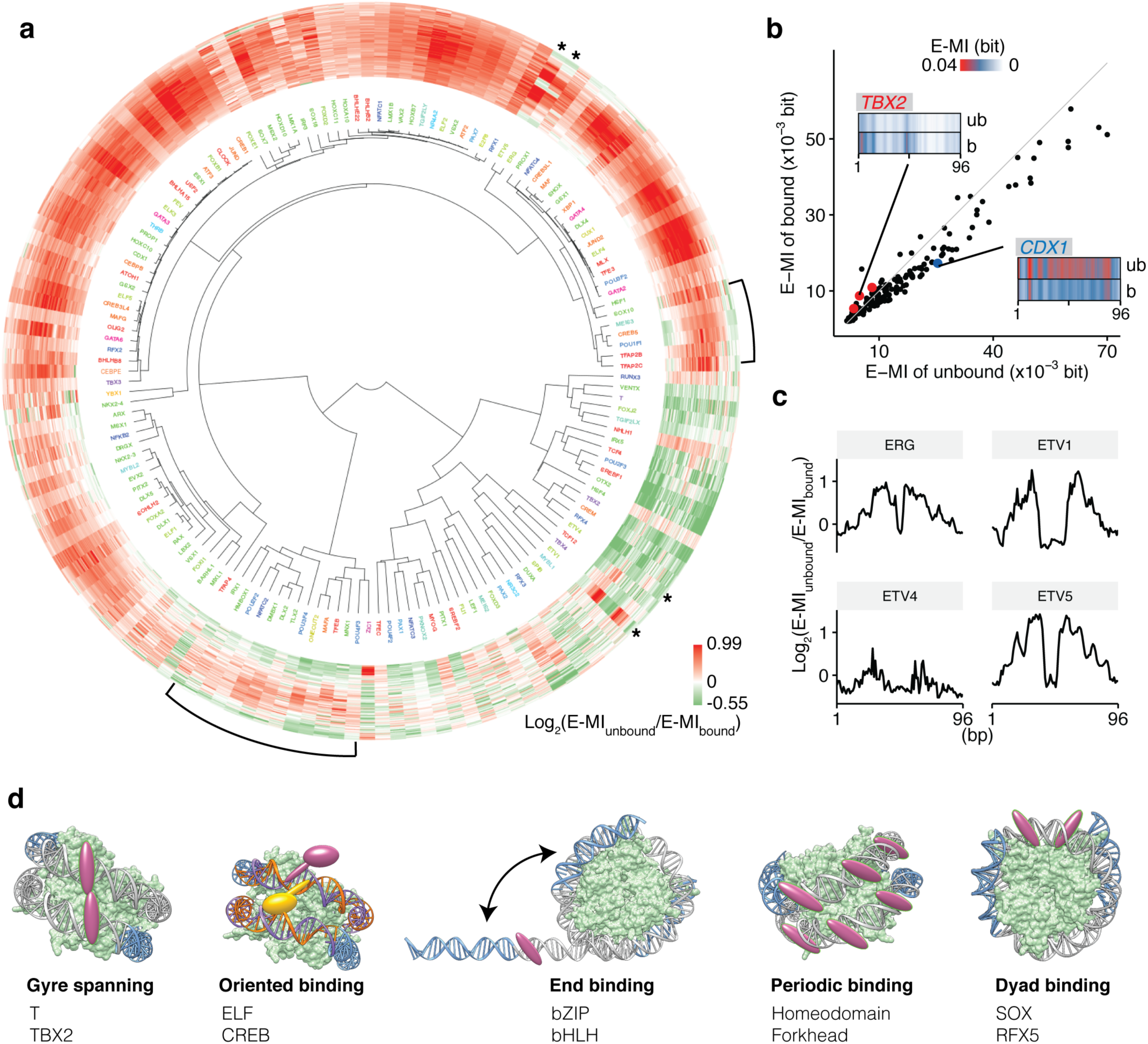
Effects of TF binding on nucleosome stability and a summary of identified TF-nucleosome interaction modes. **a**, Hierarchical clustering of the differential E-MI diagonal between the bound and the unbound cycle 5 libraries. TF names are colored to encode their family information (coloring scheme as indicated in **Fig. 2a)**. Brackets denote TFs that both destabilize and stabilize nucleosome in a position-dependent way. Asterisks denote the ETS factors with a specific pattern of positional dependence. **b**, Mean strengths of E-MI diagonals in the bound and the unbound cycle 5 libraries. The scatterplot shows the mean E-MI for the diagonals of each TF (dots), and for both the bound library (*y* axis) and the unbound library (*x* axis). The grey line represents where *y=x*. Most TFs have stronger signals in the unbound library (e.g. CDX1, blue). A few TFs show the reverse (e.g. TBX2, red). For CDX1 and TBX2 the E-MI diagonals of the bound (b) and the unbound (ub) libraries are also illustrated. **c**, Differential E-MI diagonals for the four ETS family TFs indicated by asterisks in **(a). d**, The identified major interaction modes of TFs with nucleosomal DNA.

We also found several cases where different binding modes of the same TF could dissociate nucleosome with a different efficiency (**Extended Data Fig. 8b**). Moreover, many TFs’ efficiency to dissociate nucleosome depended on the position of binding. In general, we observed that binding events close to the center of nucleosomal DNA more efficiently dissociated the nucleosome (**Fig. 7a** and **Extended Data Fig. 8a**). Interestingly, some TFs could both stabilize and destabilize nucleosome in a position-dependent way. Most of them tend to facilitate the dissociation of nucleosome when bound close to the center of the nucleosomal DNA, and stabilize the nucleosome when bound to the ends (**Fig. 7a**, brackets). It is possible that TFs bound close to the ends could decrease the DNA flexibility there and subsequently disfavor the dissociation of DNA ends from the histones, which in turn contributes to nucleosome stability. More specifically, some ETS members decrease in their efficiency to dissociate nucleosome or even stabilize nucleosome when they bind very close to the dyad (e.g. the ETV factors and ERG, asterisks in **Fig. 7a**, see also **Fig. 7c**).

## DISCUSSION

It is well established that TFs compete with nucleosomes for available genomic DNA sequences, and that this competition has a major influence on gene expression. Although the DNA binding specificities of many TFs and the nucleosome itself are relatively well characterized^38,42,43,53-60^, there is little information on how the nucleosome affects TF binding. In this study, we developed a new method, NCAP-SELEX, for analysis of nucleosome-TF interactions and systematically examined 220 TFs’ binding preference on nucleosomal DNA. To identify the binding patterns, we used a mutual-information-based method that can detect enrichment of any sequence pattern along the nucleosomal DNA. This analysis, combined with motif matching, identified five major interaction patterns between TFs and the nucleosome (**Fig. 7d**). The interaction modes include (1) binding spanning both of the two gyres of nucleosomal DNA; (2) orientational preference; (3) end preference; (4) periodic binding; and (5) preferential binding to the dyad region of nucleosomal DNA. Together, these findings reveal a rich landscape of interactions between the two key regulators of genome structure and function—the nucleosome and the sequence-specific DNA binding proteins.

### Nucleosomes mask interaction surfaces on DNA

Our results confirmed the previous view^18^ that the nucleosome inhibits binding of almost all TFs to DNA. TFs and the nucleosome have long been considered to bind DNA in a mutually exclusive fashion^30,61,62^. However, only in a few individual cases has this prediction been validated using direct biochemical assays^19,63^. Here, we performed an NCAP-SELEX experiment that analyzes TF-nucleosome interactions in the absence of higher order effects, such as chromatin compaction, remodeling or histone modification, which may complicate analysis of the *in vivo* TF-nucleosome interactions. We find that for almost all TFs, less binding occurs in regions that have higher nucleosome occupancy (**Fig. 2a**). This result directly verifies the inhibitory role of the nucleosome. In addition, we observed that although differing in extent, most TFs prefer to bind nucleosomal DNA close to the entry and exit positions (**Fig. 4a**). This positional preference is in line with the probability of spontaneous dissociation (breathing) of nucleosomal DNA, which decreases from the end to the center^64-66^. Therefore, the end-binder class of TFs may only be able to bind to regions of DNA that are dissociated from the nucleosome.

Wrapping of DNA around the nucleosome results in masking of one side of the DNA helix, but leaves the other side accessible from solvent. Such masking results in a significant accessibility change along each period (~10.2 bp) of the nucleosomal DNA. Therefore, the nucleosome will directly sterically hinder TFs that bind to long motifs through a continuous interaction with the major or minor groove. In particular, this could block the binding of the C2H2 zinc fingers (**Fig. 4a**, see also the references^26,54,55^). In addition, strong steric hindrance will block binding of proteins that radially cover more than 180° of the DNA circumference. This may explain the observed end preference of, for example, the bZIP family and many bHLH factors (**Fig. 4**, see also the references^26,67^). Moreover, nucleosomal DNA is bent relatively sharply, which could impair TF-DNA contacts if TFs have evolved to specifically bind to free DNA.

However, we found that many TFs that bind to short motifs, or to discontinuous motifs, are still able to bind to nucleosomal DNA in a periodic pattern that corresponds to the helical periodicity of DNA. This periodicity was not observed on free DNA, indicating that the occlusion of specific positions by the nucleosome still allows TFs to occupy the remaining sites. Such periodic preference of binding has been reported previously for p53 and the glucocorticoid receptor^68,69^, but the prevalence and biochemical basis of this phenomenon was not clear.

### Nucleosome leads to asymmetric binding of TFs

Our analysis identified many TFs such as ETS and CREB that have an orientational preference to nucleosome when binding nucleosomal DNA. The asymmetry is also observed for the MNase (**Fig. 3f, Extended Data Fig. 3e**) and DNase I^70^ profiles around their *in vivo* binding sites. Such orientational preference is induced by the nucleosome, because the nucleosomal environment breaks the rotational symmetry of DNA. Asymmetric chromatin features have been extensively observed previously by many investigators. These include signatures like nucleosome occupancy^71^, chromatin accessibility^70^, histone modification^71,72^, and the nucleosome signatures^73,74^. As these features are a complex outcome of many active and passive cellular processes, the origin of the observed polarity has been unclear. Our results suggest that at least part of the observed asymmetry in chromatin features next to TF binding sites or across nucleosomes is the direct result of the fact that TFs can interact with the nucleosome in a preferred orientation. In addition, because many TFs, including canonical homeodomains, recognize a near-palindromic site even when they bind DNA asymmetrically, the orientational asymmetry is likely to be more pervasive than what was detected in this study.

### Nucleosome as a scaffold

The nucleosome has DNA wrapped around it and acts as a scaffold, facilitating specific binding modes that would display very weak affinity on free DNA. A unique property of the nucleosomal DNA is that at most positions, two DNA gyres are parallel to each other. Moreover, the DNA grooves align across the two nucleosomal DNA gyres^41^. The parallel gyres could specifically associate with TF dimers, or TFs having long recognition helices or multiple DNA binding domains. Here, we found the T-box factors T and TBX2 are using this scaffold to bind nucleosomal DNA. Similar multi-gyre binding has previously been reported for synthetic DNA binder^75^ and for large protein complexes involved in chromatin remodeling^76,77^. However, our results are the first demonstration of this mode of binding for sequence-specific DNA binding proteins. The dual-gyre binding is possible only on nucleosomal DNA, and it thus stabilizes the nucleosome from dissociation, and may therefore function to lock a nucleosome in place at a specific position.

In addition to the dual-gyre binding mode, we also identified several TFs that prefer to bind at or near the dyad axis. These included RFX5 and five SOX TFs. The dyad region of nucleosomal DNA differs from other nucleosomal DNA in three respects. First, the dyad region contains only a single DNA gyre and thus has a lower steric barrier for binding. Second, the histone disk of the nucleosome is thinnest near the dyad; this further reduces the steric barrier, and also allows TFs to deform the dyad DNA more easily due to a weaker interaction with histones; the higher deformability likely accounts for the dyad preference of SOXs, which bend DNA upon binding. Third, the entry and exit of nucleosomal DNA are also close to the dyad; together with the dyad DNA, they provide a scaffold for specific configurations of TFs. FoxA has been suggested to make use of this scaffold to achieve highly specific positioning close to the dyad^20,78^; this binding mode mimics that of the linker histones H1 and H5^79^. However, the dyad positioning of FoxA is not observed in this study using eDBD, potentially because the full length of FoxA is required for its interaction with the nucleosome^21^.

Available sites on histones also contribute to part of the nucleosome scaffold. Many proteins bind nucleosomal DNA by contacting both the nucleosomal DNA and the histones, as evidenced for the chromatin remodelers and histone modifiers^51,80,81^. The additional contact with histones will allow proteins to bind nucleosomal DNA with a higher affinity than free DNA, and could also lead to functional histone distortions upon binding^82^. Further structural analyses are necessary to determine whether the positional preferences of, for example, SOXs and RFX5 are resulted from interactions with histone proteins, or are primarily driven by the more accessible nature of the dyad DNA.

### Pioneer TFs and nucleosome binding

Pioneer TFs are defined by their ability to bind nucleosomal DNA^18^. In many cases it is unclear whether such TFs prefer nucleosomal DNA over free DNA, or bind nucleosomal DNA only relatively better than non-pioneer TFs. It is noteworthy that TFs may also facilitate the access of nucleosomal DNA even without displacing the nucleosome, by competing with linker histones and maintaining nucleosome in an accessible conformation^20^. Such a mechanism requires a dyad preference. It is thus of particular interest to further examine the interaction of dyad binders with linker histones.

In NCAP-SELEX, we observed that for the eDBD of almost all TFs, including known pioneer factors such as FOX and SOX, binding to free DNA was preferred compared with their binding to nucleosomal DNA. This order of preference results in destabilization of the nucleosomes that are bound by the TFs by mass action. The different binding modes that we identified also differ in their potential for pioneer activity. Whereas end-binders are unable to effectively access nucleosomal DNA, a large fraction of nucleosome-bound DNA sequence will be accessible to the TFs in the periodic binder class. The dyad binders, in turn, can access only highly specific positions along the nucleosomal DNA. Moreover, some TFs have developed “pioneer modes” to bind nucleosomal DNA in a different way compared to their binding to free DNA. For example, the transcription factor T has its normal binding mode inhibited by nucleosome occupancy (**Fig. 3a**, **Extended Data Fig. 2a**). Nonetheless, its dual-gyre binding mode is only allowed on nucleosomal DNA. It is also possible that we did not identify some pioneer TFs, as additional domains in the full-length protein could be required for their high-affinity binding to the nucleosomal DNA. The ability of a large number of TFs to access different positions along the nucleosomal DNA indicates that nucleosomes at different genomic positions are accessible to different classes of TFs, leading to a complex interplay between the DNA sequence, nucleosome positions, and the TF content of a cell.

### Dissociating the nucleosome

The binding of pioneer factors does not necessarily dissociate the nucleosome. But their ability to dissociate nucleosomes is linked to their tendency to open chromatin and to activate transcription. For the libraries enriched by each TF, we examined if the nucleosome is dissociated by comparing the bound and unbound libraries of cycle five. In accord with the mutually exclusive nature between TF and nucleosome binding, most TFs facilitated the dissociation of nucleosomes. In cells, these TFs are predicted to act passively to dissociate nucleosomes, by having a moderate affinity towards nucleosomal DNA, and high affinity towards free DNA. This mechanism provides favorable kinetics as binding would not require prior dissociation of the nucleosome, and also contributes free energy for displacing the nucleosome. These TFs are thereby potential activators that can open chromatin and regulate gene expression.

Some TFs, in turn, stabilized the nucleosome. These factors could act to repress gene expression, or to precisely position nucleosomes at specific genomic loci. However, in cells, they might also potentially activate an enzymatic process that leads to dissociation, displacement or remodeling of the nucleosome. Moreover, we also observed TFs that both stabilize and destabilize nucleosomal DNA depending on their relative position of binding. Such ability could be used to more precisely position local nucleosomes.

TFs and the nucleosome are central elements regulating eukaryotic gene expression. In this work, we have systematically analyzed the ability of TFs to bind to and to dissociate the nucleosome. The results revealed five distinct modes of TF-nucleosome interactions, including a symmetry-breaking effect induced by the nucleosomal context that is likely to contribute to the extensively observed asymmetric environment around gene regulatory elements. In addition, we discovered major differences in the ability of specific TFs to bind to and open nucleosomal DNA. The identified binding modes explain in part the complexity of the relationship between sequence and gene expression in eukaryotes, and provide a basis for future studies aimed at understanding transcriptional regulation based on biochemical principles.

## END NOTES

**Supplementary Information** is linked to the online version of the paper.

## Acknowledgments

We thank Biswajyoti Sahu, Fan Zhong, Kazuhiro Nitta, Arttu Jolma, Minna Taipale, Bei Wei, Jilin Zhang, Ekaterina Morgunova, and Jarkko Toivonen for their valuable suggestions. We thank Lijuan Hu, Jianping Liu, and Sandra Augsten for technical assistance. We thank the ENCODE Consortium and the ENCODE production laboratories for generating the ChIP-seq datasets. This work was supported by Center for Innovative Medicine at Karolinska Institutet, Knut and Alice Wallenberg Foundation, Göran Gustafsson Foundation and Vetenskåpsradet. P.C. was supported by the Deutsche Forschungsgemeinschaft (SFBB60, SPP1935), the European Research Council Advanced Investigator Grant TRANSREGULON (grant agreement No 693023), and the Volkswagen Foundation.

## Author Contributions

J.T., F.Z. and P.C. conceived the experiments. F.Z. performed the NCAP-SELEX, MNase-seq and data analysis. L.F produced the histone octamers. Y.Y. helped with the curation and production of TF proteins. E.K. analysed the MNase-seq data. S.D. performed EMSA validation for SOX. F.J. and J.T. interpreted the data and wrote the manuscript. All authors discussed the findings and contributed to the manuscript.

## Author Information

All next generation sequencing data will be deposited to European Nucleotide Archive (ENA) under Accession PRJEB22684. All computer programs and scripts used are either published or available upon request. The authors declare no competing financial interests. Requests for materials should be addressed to J.T.

## METHODS

### Preparation of histone octamer

A vector encoding *Xenopus laevis* H2A with an N-terminal tag was cloned using ‘Round-the-horn site-directed mutagenesis. *X. laevis* histones were expressed and purified as described previously^39^. Inclusion bodies were resuspended by using a Dounce tissue grinder (Sigma-Aldrich). Purified histones were aliquoted, flash-frozen, lyophilized, and stored at -80°C prior to use. The lyophilized histones were resuspended in unfolding buffer (7 M guanidine hydrochloride and 10 mM DTT in 20 mM Tris-Cl, pH 7.5) to a concentration of 1.5 mg/ml. N-terminally tagged H2A, H2B, H3 and H4 were then combined at a molar ratio of 1.2:1.2:1:1. The sample was incubated on ice for 30 minutes before it was dialyzed against three times 600 ml refolding buffer (2 M NaCl, 1 mM EDTA, and 5 mM β-mercaptoethanol in 10 mM Tris-Cl, pH 7.5). The sample was recovered after dialysis and applied to a GE S200 16/600 pg size exclusion column (GE Healthcare, Little Chalfont, United Kingdom). Peak fractions were analyzed by SDS-PAGE. Fractions containing the octamer were pooled and concentrated. Both the histone expression and octamer formation have been quality-controlled (**Extended Data Fig. 1a, b**).

### Clones, protein expression and purification for TFs

Gateway recipient vectors having a pETG20A backbone were employed in the bacterial protein expression. Insertions for these expression vectors were derived either from PCR clones or from gene synthesis; the details are given by Yin *et al*.^54^. The sequences and domains for all TFs are listed in **Supplementary Table 1**. The non-full-length constructs contain extended DNA binding domains (eDBDs), with a design rationale reported previously^38^. We essentially followed Yin *et al.^54^* to express and purify the proteins from *E. coli* cells.

### Nucleosome CAP-SELEX

The Nucleosome CAP-SELEX (NCAP-SELEX) protocol has two steps of selection, respectively for ligands bound by the nucleosome and by individual TFs. The DNA ligands were designed based on Illumina’s Truseq library (**Supplementary Table 2**). The adapter lengths were 24 bp at the left side and 22 bp at the right side. The total lengths of the ligands are 147 bp for lig147, and 200 bp for lig200, with 101 bp and 154 bp in random, respectively. The single-stranded oligos of lig147 and lig200 were purchased from IDT (Ultramer DNA oligos). A PCR reaction with primers binding to the adapters (**Supplementary Table 2**, PCR_primers) was used to obtain double-stranded DNA from the synthetic oligos, and was also used to amplify the libraries between SELEX cycles. For sequencing, the ligands were amplified with the multiplexing primers (**Supplementary Table 2**, PE_PCR_primers).

In SELEX, first, double-stranded DNA ligand and tagged histone octamer were mixed in 2 M KCl solution and incubated for 30 min. The mixture was then diluted stepwise as described by Dyer *et al.^39^*, with a dilution buffer (TE buffer supplemented with 1 mM tris(2-carboxyethyl)phosphine (TCEP) and a cocktail of protease inhibitors (05892970001, Roche)). The reconstituted nucleosome was incubated for 30 min with the corresponding affinity beads (pre-blocked with the blocking buffer containing 25mM Tris, 0.5% BSA, 0.1% tween 20, 0.02% NaN_3_), and at the same time shaken under 1900 rpm with a microplate shaker (13500–890, VWR). The beads were then washed 15 times with a microplate washer (Hydrospeed™, Tecan). The nucleosome was eluted and incubated with 10–200 ng purified TFs for 20 min. The TF-bound species were pulled down with magnetic beads (pre-blocked with the blocking buffer) and washed 15 times. The bead suspension were used for PCR as previously described by Jolma *et al*.^38^ This process was repeated for four cycles. Ligands were amplified and sequenced after each cycle and before the experiment (the input). When incubating the nucleosome with TF, we initially used 140 mM of monovalent cations. The physiological salt concentration resulted in relatively high nonspecific adsorption of the nucleosome to the sepharose beads. To improve the assay, lower salt concentrations (50 mM to 75 mM) were used in subsequent experiments. Most effects were robust to the changes in the salt concentration; discussion in the main text is limited to observations that were detected under multiple salt concentrations. Moreover, in SELEX, each cycle is essentially an independent replicate of the experiment. The reported effects all show enrichment across multiple SELEX cycles.

To interrogate whether the binding of TFs facilitates the dissociation of nucleosome, we carried out a fifth cycle that further separated the TF-bound species into libraries unbound and bound by nucleosome. The TF-bound species were depleted for the nucleosome-bound species with affinity beads for the tag on the histone. The ligands bound by the beads were collected as the nucleosome-bound libraries. The DNA ligands remaining in the supernatant were sequenced as the unbound libraries. As a control, the cycle five nucleosome was also allowed to dissociate in the absence of TFs; the bound library and the unbound library were collected as described above.

As a control, HT-SELEX (SELEX using free DNA) with lig147 or with lig200 was performed according to the previous protocol^54,55^ with the same purified TF proteins as those used in NCAP-SELEX.

The input amount of DNA will exhaust almost all possible 20-bp consecutive or gapped subsequences. Such complexity well suffices the specificity studies of human TFs, whose binding is associated with ~15 bits of infomation on average^55^. For nucleosome, the complexity allows the study of optimal sequences around each histone-DNA contact, but might not capture all the specificities as the nucleosome-favored or disfavored sequences may include cooperation spanning a large length of DNA, e.g., the phased successive bending or the rigidity of a long segment.

The NCAP-SELEX and HT-SELEX library for each TF contains hundreds of thousands of unique reads. Under this sample size, if a TF is binding nucleosomal DNA without restrictions, any non-random pattern of TF binding that has a biologically meaningful effect size (as observed in our study) can only occur with an extremely small p-value.

### Sequencing and pre-processing

The SELEX ligands amplified with multiplexing primers were purified with AMPure beads (Beckman Coulter), and sequenced using Illumina Hiseq 2000 or Hiseq 4000, with >80 bp paired-end settings. Raw sequences were demultiplexed with bcl2fastq (v2.16.0.10). In general hundreds of thousands of reads were obtained for each TF.

The R1 and R2 reads of paired-end sequencing were merged with PEAR^84^ requiring 5 bp overlap at minimum. The merged sequences were discarded if their variable region length is not the same as the ligand design. The obtained sequences were then trimmed for adaptor and for quality by Trim Galore (version 0.4.3). All shorter sequences produced during trimming were removed. The sequences were further cleaned for adaptor sequences by removing all sequences that contained a 14-bp overlap with Illumina sequences. For further analysis, we removed the PCR duplicates and used only the unique reads.

### TF signal analysis with E-MI

MI between the most enriched 3-mer pairs (E-MI) was calculated for all non-overlapping position combinations:

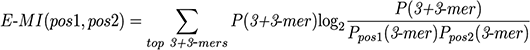
where *P(3+3-mer)* is the observed probability of a 3-mer pair (i.e. gapped or ungapped 6 mer) from position 1 and position 2. *P_pos1_(3-mer)* and *P_pos2_(3-mer)*, respectively, are the marginal probabilities of the constitutive 3-mers at position 1 and position 2. Their product represents the expected probability of the 3-mer pair. Sums are over the top 10 most enriched 3-mer pairs, which have the highest ratio between the observed probability and the expected probability. For the diagonal plot, E-MIs from position pairs where *pos2* = 3 + *pos1* were used.

Clustering of the E-MI diagonal was performed using the cosine distance metric and ward.D2 linkage of the *hclust* function in R. The circular representation of the classification result was generated using the *circlize* R package^85^. To calculate the penetration of E-MI for each TF, the diagonal of E-MI was first LOESS smoothed with a span of 0.45; next, for each half of the diagonal, the maximum E-MI value among the half was identified; after that, the positions where the E-MI decreases to half of the E-MI maximum were taken as the penetration depth. The final penetration depth is the average value for both halves of the E-MI diagonal.

To check whether the gyre-spanning mode of TF T is preferring nucleosomal DNA, for both its bound and unbound libraries of cycle 5, the E-MI strength of Mode 2 binding was evaluated by summing E-MI from 3-mer pairs spaced 77–83 bp, the E-MI strength of the background was evaluated by summing E-MI from 3-mer pairs spaced 50–70 bp. For both the binding signal and the background, Log2 ratios of E-MI strength between the bound and unbound libraries were calculated for four independent replicates of NCAP-SELEX using TF T. The obtained ratio indicates whether the signal (or the background) has a different strength between the two libraries.

When comparing E-MI between the bound and the unbound libraries from cycle five, only TFs with the 3×FLAG tag were considered.

### Motif matching and PWM (positional weight matrix) generation

Motif matching for each TF was conducted using MOODS^86,87^ with p-value set to 0.0001. The motifs used in matching were from our previous curations^54,55^. Motif hits from both strands were combined unless indicated. When necessary, motifs from NCAP-SELEX were generated using Autoseed^38,88^ with multinomial of 1.

### Quality control of the SELEX experiments

The successful TFs were called by manually checking the E-MI and motif discovery results for each TF. The successful TFs has detectably stronger E-MI between neighboring 3-mer pairs than that between 3-mer pairs far away from each other, and show enriched motifs that are not contaminations from unrelated TFs.

### Evaluation of nucleosome-induced orientational preference of TFs

On free DNA, motifs have the same affinity for TF-binding irrespective of its orientation. This is not true when DNA is wrapped onto a nucleosome as the nucleosome breaks DNA’s 2-fold rotational symmetry. Depending on motif’s relative orientation to nucleosome, the same motifs can differ in affinity. This orientational asymmetry was examined systematically using the lig200 NCAP-SELEX libraries. For each TF, we first calculated the binding energy difference (ΔΔ*G*) between the two relative orientations for each of the most enriched non-palindromic 8-mers (top 40 used). The ligands in this TF’s SELEX library were divided into two halves according to the dyad position. The two halves were calculated separately and then averaged. Similarly to previous studies^89,90^, we assumed a low TF concentration and that the dissociation during wash is insignificant for high-affinity 8-mers. Consequently, for each 8-mer and for each half of the ligands, the ΔΔ*G* of the 8-mer between the two relative orientations is

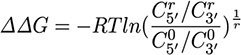

Here *C_5’_* and *C_3’_* are counts of this 8-mer, respectively for the DNA-strands with their free ends located at the 5’ and the 3’ (the other end is at the dyad where we divide). The count ratio *C_5’_/C_3’_* in cycle *r* was normalized with the count ratio in cycle 0, taken the *r*th root to account for the exponential enrichment in SELEX, and subsequently converted into energy difference. The directional energy difference for each 8-mer was then averaged for the two halves of the ligands, and the absolute value is used to represent the orientational asymmetry of this 8-mer

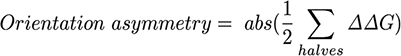

Orientational asymmetry of the TF is then represented by the mean of the 40 most enriched 8-mers’ orientation asymmetry.

To rule out any potential orientational bias induced by the adaptors of the SELEX ligands, we also calculated the orientation asymmetry values for 8-mers in the HT-SELEX library. For each TF, the 8-mers used for its HT-SELEX library are the same 8-mers as used for its NCAP-SELEX library. After obtaining the 8-mers’ orientation asymmetry values for both the NCAP-SELEX library and the HT-SELEX library of the TF, we used a one-tailed t-test to examine if the orientation asymmetry values in the NCAP-SELEX library are larger than those in the same TF’s HT-SELEX library, and obtained the p-value.

Signal enrichment in each TF’s library was represented using the median fold change of the 8-mers that are most enriched (top 10 8-mers). The fold change for each 8-mer was calculated using log_2_(cycle 4 count / cycle 0 count).

### MNase-seq

In MNase-seq, the LoVo cell line from ATCC was used (CCL-229, tested to be free of mycoplasma infection by Hoechst staining). MNase-seq was performed similarly as described previously^91^. Specifically, 10^7^ cells were harvested and washed twice with 10 ml cold DPBS (Dulbecco’s phosphate-buffered saline), spinned down with 350 g for 5 min at 4 °C. The cells were next crosslinked with 10 ml of 1.1% formaldehyde for 10 min in DPBS, tumbling end over end. The crosslinking reaction was quenched with 50 μl 2.5 M glycine and further tumbled for 2 min, and washed twice with cold DPBS. Lysis of the cells was performed with 20 ml 0.5× PBS containing 0.5% Triton X-100 for 3 min on ice; the nuclei were then collected by centrifugation (350 g, 5 min). Before MNase digestion, the nuclei were washed three times with 1 × MNase digestion buffer, resuspended with 1 ml of the same buffer containing 100 μg/ml RNase A. An aliquot of 100 μl was used for MNase digestion. MNase digestion was carried out with 100 units of MNase (M0247S, NEB) at 37 °C for 8 min, quenched with 100 μl stop buffer (40 mM EDTA, 40 mM EGTA, 1% SDS, 1.5 mg/ml proteinase K) at 65 °C o/n. The MNase fragments with length of 100–1000 bp were selected using Ampure beads (Beckman Coulter), and subjected to the library preparation workflow of Illumina (E7370L, NEB). The paired-end sequencing (2 × 86 bp) was performed using Illumina HiSeq 4000.

MNase-sequencing data from K562 cell line were downloaded from GEO accession GSE78984. Three titration series (20.6U, 79.2U and 304U) of MNase were selected.

### Combined analysis of MNase-seq and ChIP-seq

For MNase-seq data, the raw sequencing reads were quality and adapter trimmed with cutadapt version 1.12 in Trim Galore (version 0.4.3). Low-quality ends trimming was done using Phred score cutoff 30. Adapter trimming was performed using the first 13 bp of the standard Illumina paired-end adapters with default parameters. Raw sequencing reads were mapped to the human reference genome (hg19) using bwa^92^ with default parameters. Duplicates were removed with samtools (v 1.3.1) rmdup function. Insert size distribution was calculated based on 10000 reads that were aligned to autosomes. After duplicate removal, data from K562 titration series were merged.

Coverage of MNase fragments with length >140bp was calculated at ChIP-seq peaks of 20 TFs in K562 cell lines. We selected 500 highest signal ChIP-seq peaks that had respective TF’s motif match site and did not overlap with hg19 blacklist genomic regions. BEDtools (v2.26.0) genomecov and intersect functions were utilized in calculations. ENCODE narrowPeak calls including two replicates were used from March 2012 freeze (UCSC wgEncodeAwgTfbsUniform track) release for ATF3, CEBPB, CTCF, ELF1, GATA2, JUND, SRF, USF2 and YY1, and from later releases (the ENCODE Portal http://www.encodeproject.org, accessed 07/12/2017) for ATF2, CREB3L1, CREM, ELF4, HMBOX1, MYBL2, NFATC3, PKNOX1, RFX1, SREBF1 and YBX1^72^. Genomic sites recognized by each motif retrieved from previous HT-SELEX runs were searched from the human genome using program MOODS^86^ with a p-value cut-off of 10^-4^ and a score cut-off of 5. Final MNase fragment coverage values were calculated by taking the average of multiple motifs for each TF, and correlated with E-MI penetration values with Pearson’s method.

V-plots were generated as described by Henikoff *et al.^45^*. MNase-fragments aligned to autosomes were used. The LoVo ChIP-seq data from Yan et al.^37^ were downloaded from GEO accession GSM1239499 and GSM1208610. The peak calls were transformed from hg18 to hg19 coordinates by using UCSC liftOver and peaks within hg19 blacklist genomic regions were excluded. Genomic sites recognized by each motif were searched from the human genome using MOODS^86^ with a p-value cut-off of 10^-4^ and a score cut-off of 5. Center-point coordinates of MNase-fragments and motif sites within ChIP-peaks were compared using BEDtools (v2.26.0) closest function using strand information of each motif match.

### Fast Fourier Transformation (FFT) analysis and structure alignment

The diagonal of E-MI for each TF’s library was subtracted with the mean, windowed by Welch’s function, and then subjected to FFT. The obtained power spectrum was further divided with the mean of the E-MI diagonal and the length of the diagonal. We next calculated FFT-AUC (area under the curve) from the power spectrum and used it as an indicator for the ~10 bp periodicity induced by nucleosome. The FFT-AUC was calculated for frequencies ranging from 0.08–0.12 bp^-1^ and subtracted with the baseline level (estimated from 0.14–0.3 bp^-1^). The phase of FFT was examined at 0.102 bp^-1^. The same process was applied to the TA dinucleotide counts across all positions of the ligand for the NCAP-SELEX library of all individual TFs.

To mimic the in-phase and out-of-phase bindings of the periodic binders with respect to the preferred TA positions on nucleosome, the available structure of TF-DNA complex was aligned to the nucleosome by matching the center of the TF’s core binding sequence either to the TA step (in phase), or to a step 5-bp downstream of the TA step (out of phase). The 6-bp core binding sequence in the structure of TF-DNA complex is defined according to the most enriched 6-mers in this TF’s NCAP-SELEX library. To make the alignment, C1–C4 on all deoxyribose rings were matched between the 6-bp core binding sequence and the 6-bp nucleosomal DNA centered in-phase or out-of-phase to the TA step.

### Electrophoretic mobility shift assay (EMSA)

Nucleosomes were formed essentially as described previously^39^ from the histone octamers and the modified Widom 601^93^ DNA sequence CTGGAGAATCCCGGTCTGCAGGCCGCTCAATTGGTCGTAGACAGCTCTAGCACCGCTTAAACGCACGTACGGTATTGTTTATTTTGTTCCTCCGCCAAGGGGATTACTCCCTAGTCTCCAGGCACGTGTCAGATATATACATCCTGT. The modified Widom 601 is embedded with a SOX11-binding segment (GGTATTGTTTATTTTGTTCCT) at the center. The sequence of the embedding segment is extracted from a ligand in the cycle 4 SELEX library of SOX11. The embedding position on Widom 601 is the same as the segment’s original position on the SELEX ligand. Nucleosomes were reconstituted using this modified Widom 601 ligand and subsequently heat-shifted at 55°C for 30 min. Next the nucleosomes (containing 1 μg DNA) were incubated on ice with purified SOX11 eDBD in a 40 μl volume. As a control, the SOX11 eDBD were also directly incubated with 1 μg modified Widom 601 ligand in 40 μl volume. The samples were then subjected to EMSA. A 0.8% agarose gel was cast and run in the 0.2x Tris–Boric acid–EDTA (TBE) buffer. EMSA was performed in native conditions at 4°C for 1 h at 120 V, and later the gel was post-stained in DNA Stain Clear G (Serva). The DNA ladder 100 bp (NEB) was used as the marker.

## EXTENDED DATA FIGURES

**Extended Data Figure 1.**
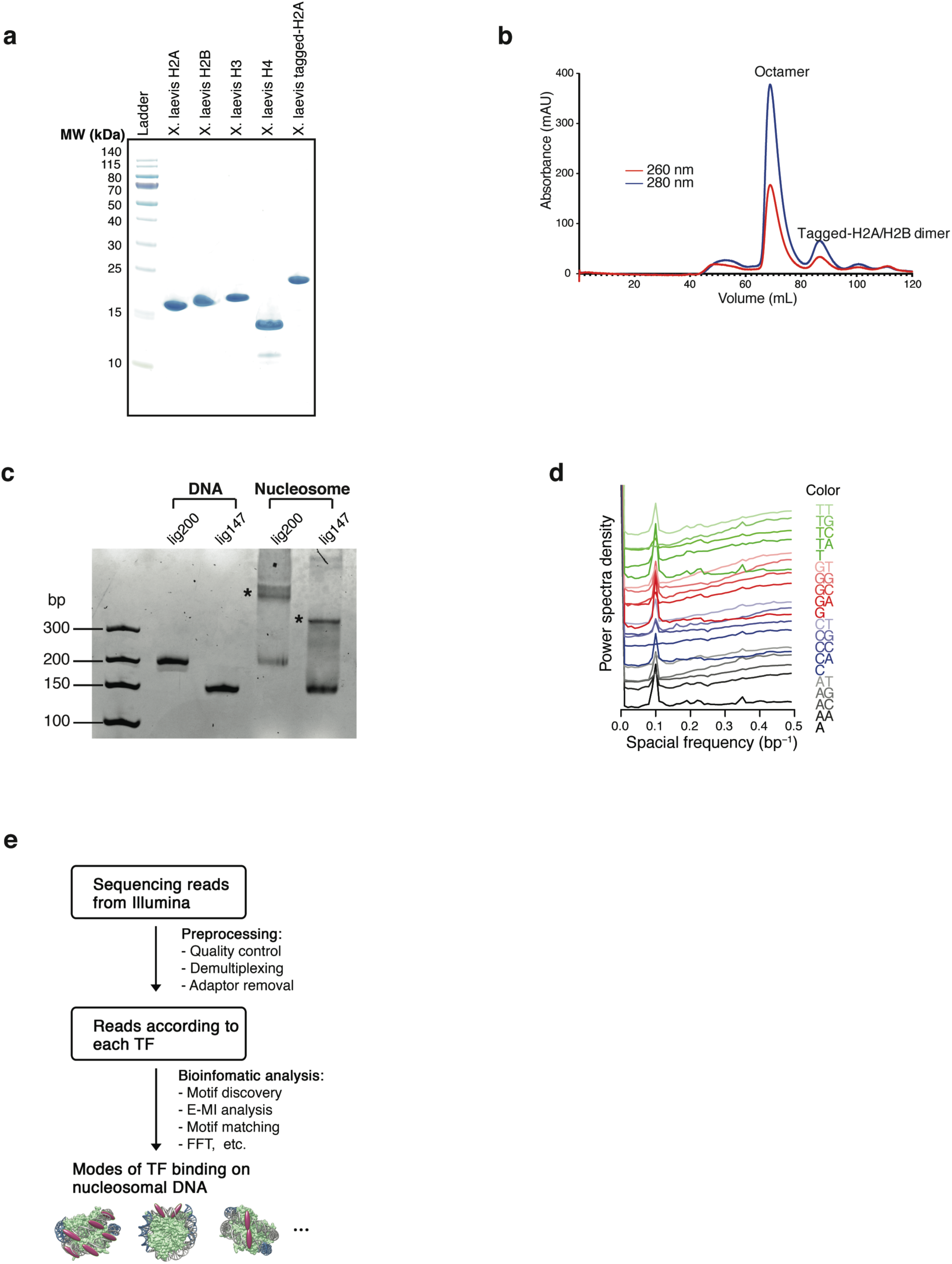
Quality control and analysis pipeline. **a**, Expression of the recombinant histones from *X. laevis*. For each lane 3 μg histone is loaded. **b**, Size-exclusion chromatogram of the histone octamer. **c**, EMSA result showing the reconstituted nucleosomes using lig147 and lig200. The original ligands are also loaded as reference. The asterisks indicate the nucleosome bands. **d**, Oligonucleotide periodicity in the library enriched by nucleosome. As a quality control of nucleosome reconstitution, we verified whether nucleosome by itself is enriching the previously reported ~10-bp periodic oligonucleotide signal^93,94^. Nucleosome SELEX (without TF) were carried out for four cycles to enrich nucleosome-favoring ligands. The counts of each single and di-nucleotide across each individual ligand were Fourier transformed and summed up for the whole library. A clear peak around 0.1 bp^-1^ (corresponding to the reported ~10-bp periodicity) is visible for most mono and dinucleotides. **e**, Analysis pipeline for the ligands enriched in NCAP-SELEX.

**Extended Data Figure 2.**
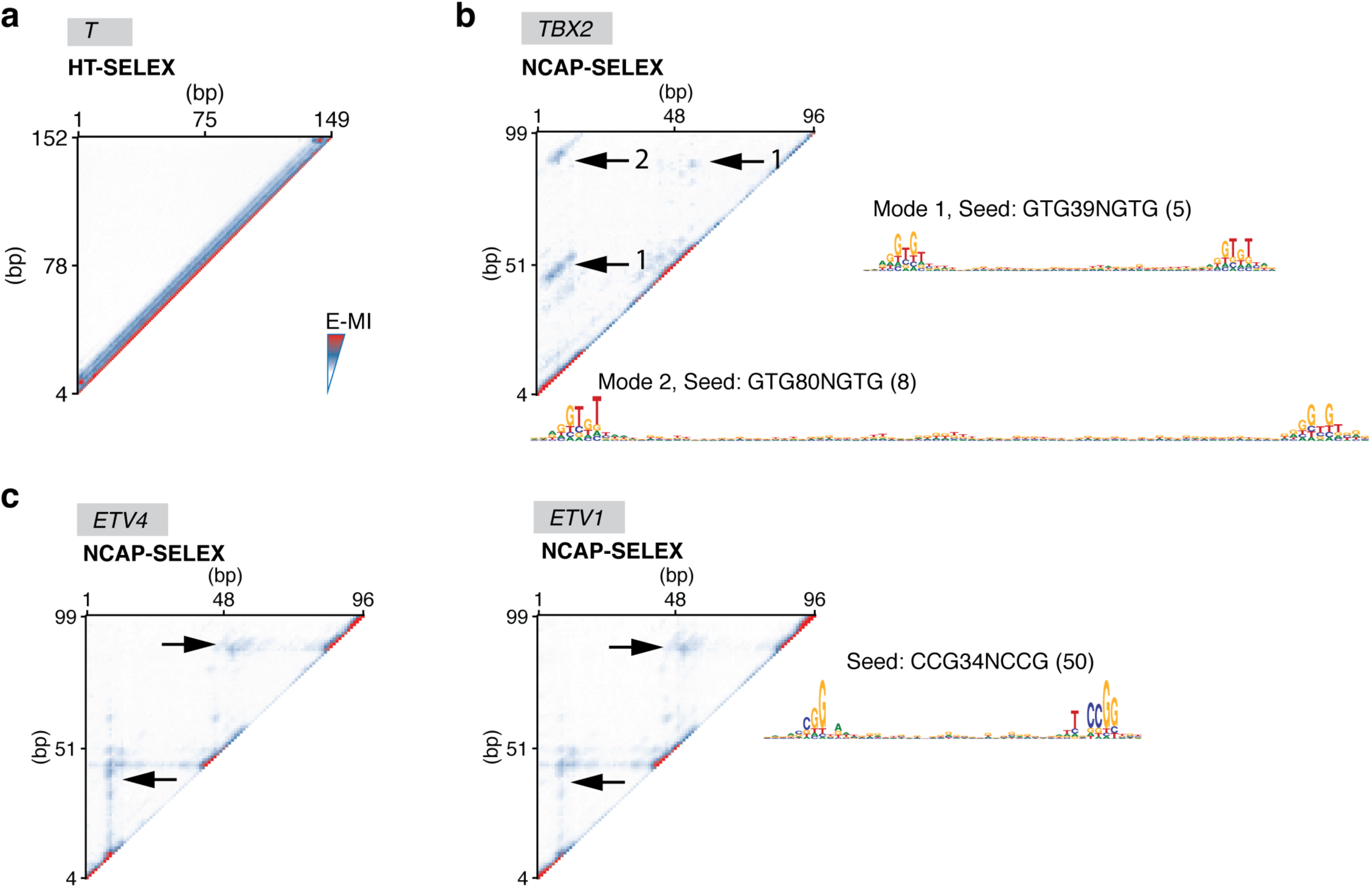
Specific binding modes allowed on nucleosome. **a**, E-MI heatmap of T (brachyury) in HT-SELEX using lig200. Pairwise E-MI for all 3-mer pairs is presented as a heatmap. The signal is only visible near the diagonal, no E-MI signal across ~80 bp is detected. **b**, E-MI heatmap of TBX2 in NCAP-SELEX using lig147. The E-MI signals across ~80 (mode 1) and ~40 bp (mode 2) are indicated. The corresponding motif of each mode is derived with the indicated seed for a specific position (number in the parentheses) in the high E-MI regions. PWM generation follows our previous method^95^ using multinomial 1. **c**, E-MI heatmap of ETV4 and ETV1 in NCAP-SELEX using lig147. The E-MI signal across ~40 bp is indicated. The motif is derived as in **(b)**.

**Extended Data Figure 3.**
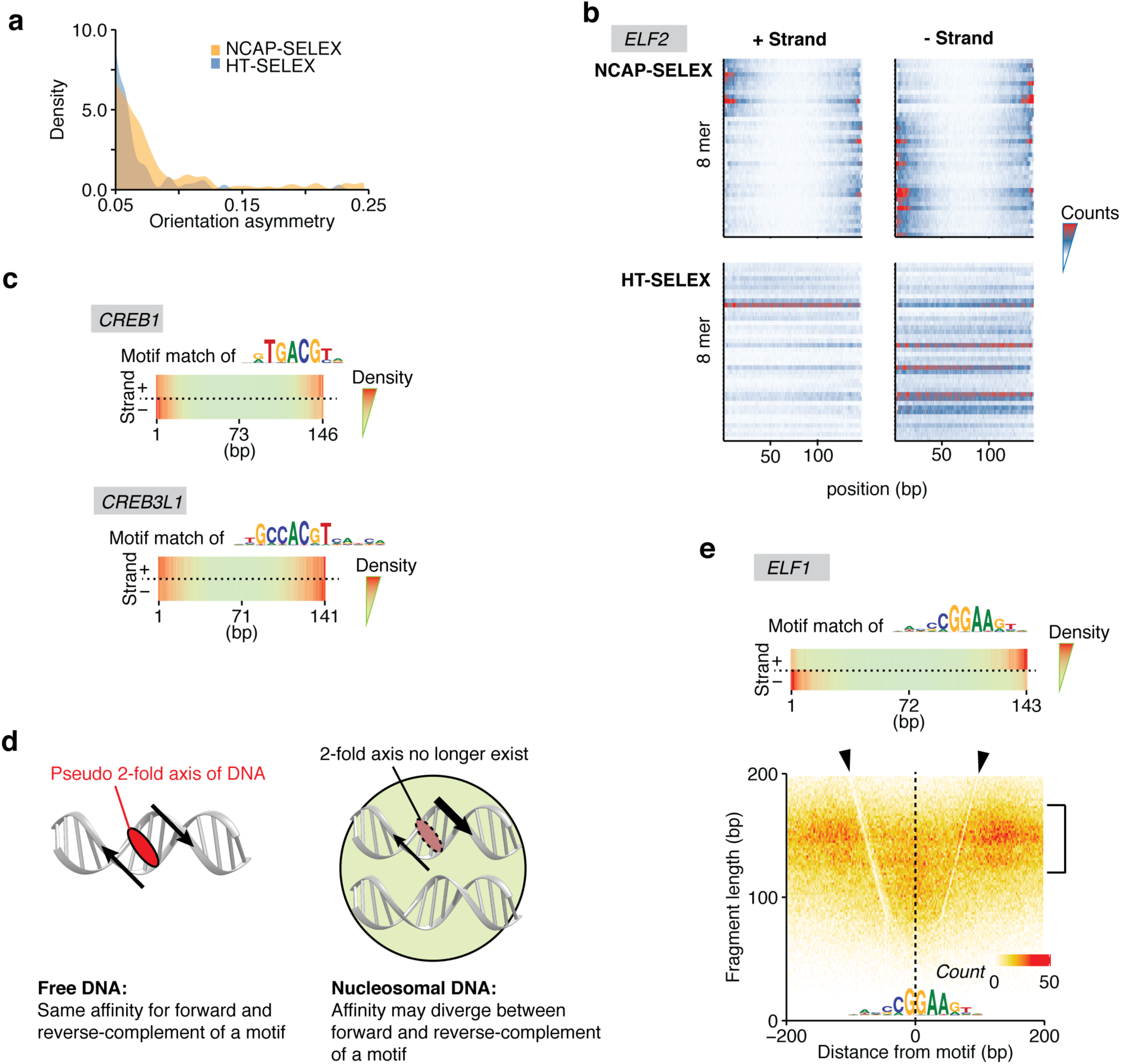
Nucleosome breaks the rotational symmetry of DNA. **a**, Density plot representing the orientation asymmetry of all TFs in NCAP-SELEX and in HT-SELEX. In NCAP-SELEX, more TFs bind with high orientational asymmetry than in HT-SELEX. A few TFs can prefer different ends of the ligand for the two binding directions in HT-SELEX; this is likely induced by the adaptor sequences. However, there are more TFs with higher orientational asymmetry in NCAP-SELEX libraries, despite the fact that for most TFs their signals are stronger in HT-SELEX libraries. **b**, Orientation asymmetry of ELF2 revealed by using top 8-mers. Each row of the heatmap corresponds to the counts distribution of a top 8-mer (non-palindromic) across the positions of the SELEX ligand. Hits of the top 8-mers occur at different ends for different strands of nucleosomal DNA (i.e. an 8-mer and its reverse-complement prefer different ends), whereas their distribution is relatively homogeneous for free DNA. **c**, Orientation asymmetry of CREB TFs. CREB TFs have different motif density distributions for the two strands of nucleosomal DNA. The motif used for matching is indicated above. The “−” strand profile is from the density of the reverse-complement motif. **d**, Break of the 2-fold rotational symmetry of DNA induces preferred orientation of TFs. Left: free DNA has a 2-fold axis (red ellipse) perpendicular to the helix axis. Motifs in two orientations are symmetric with each other with respect to a 180° rotation centered on the axis. Right: for motifs on nucleosomal DNA, if the other strand of DNA or the histone proteins (green) affects binding, the 2-fold axis of DNA no longer exists, as a 180° rotation centered on the axis no longer generates an identical conformation (the rotated image not superimposable with the original one). **e**, Orientational asymmetry of ELF1 on nucleosomal DNA. Similar to ELF2, ELF1 has different motif density distributions for the 2 strands of nucleosomal DNA (top panel). The distribution of MNase fragments around genomic ELF1 sites is also asymmetric (bottom panel); footprint of ELF1 is indicated with the arrowheads (the V-shaped lines with a lower signal density), and the range of flagment length that corresponds to nucleosome occupasion are indicated with the bracket.

**Extended Data Figure 4.**
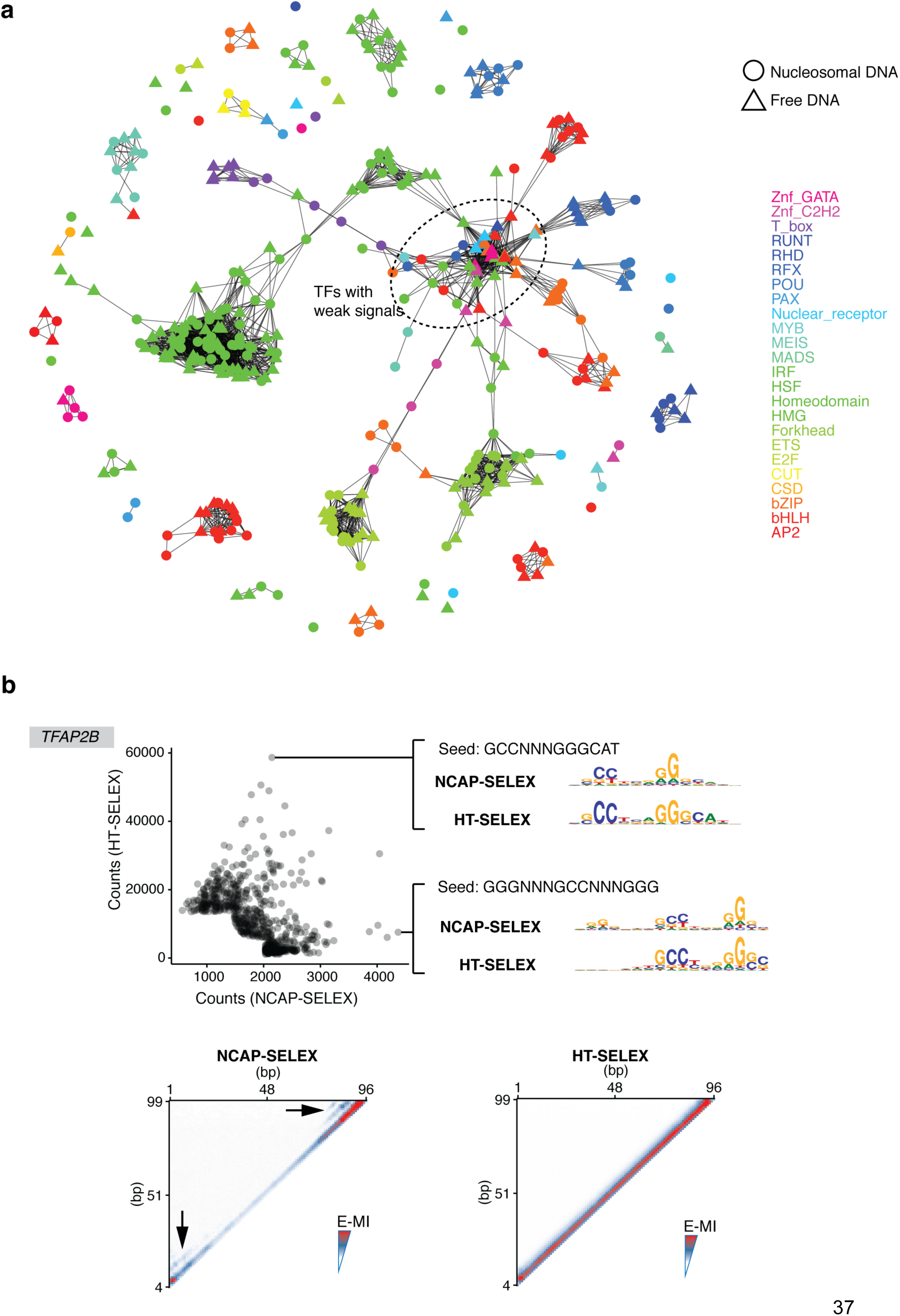
Most TFs bind nucleosomal DNA without significant motif change. **a**, Network representation of TFs’ specificities in presence and absence of the nucleosome. Vertices indicate the binding specificity profiles of TF eDBDs, either in the presence (circle) and in the absence (triangle) of the nucleosome. Vertices are colored according to the TF’s family. Two vertices are connected by an edge if the profiles they represent are similar. Specifically, we assume that all 9-mer counts in the library enriched by a TF will serve as a profile to represent its binding specificity. To evaluate the similarity between two profiles, we selected the most abundant 9-mers (top 0.1%) from either of the profiles, and calculated Pearson’s correlation using counts of these 9-mers from both of the two profiles. An edge is drawn between the profiles (vertices) if the calculated correlation is greater than 0.2. TFs from the same family generally clusters together regardless of the presence of nucleosome, indicating that TFs’ binding specificity is not significantly affected by nucleosome. TF profiles with weak signals also tend to cluster together (the cluster circled by dashed line), as the top 9-mers in their libraries are dominated by SELEX bias (e.g. the bias from PCR or wash) rather than by the TFs’ specificities. **b**, TFAP binds nucleosomal DNA with slightly different specificity than free DNA. The scatter plot (top panel) shows the counts of gapped 9-mers from SELEX libraries of TFAP2B, enriched with NCAP-SELEX (x axis) and HT-SELEX (y axis). The examined 9-mers consists of three segments of trimers interspaced with two gaps (0–5 bp). Only the most enriched 9-mers (top 300 in each library and in the combined library) are shown from clarity. For comparison, the most differentially enriched gapped 9-mers were also used as seeds to derive the corresponding motifs from both libraries (right). The heatmap (bottom panel) shows the pairwise E-MI for all combinations of positions on lig147, in the presence (left) and absence (right) of nucleosome. The arrowheads indicate the additional signals developed in the presence of nucleosome.

**Extended Data Figure 5.**
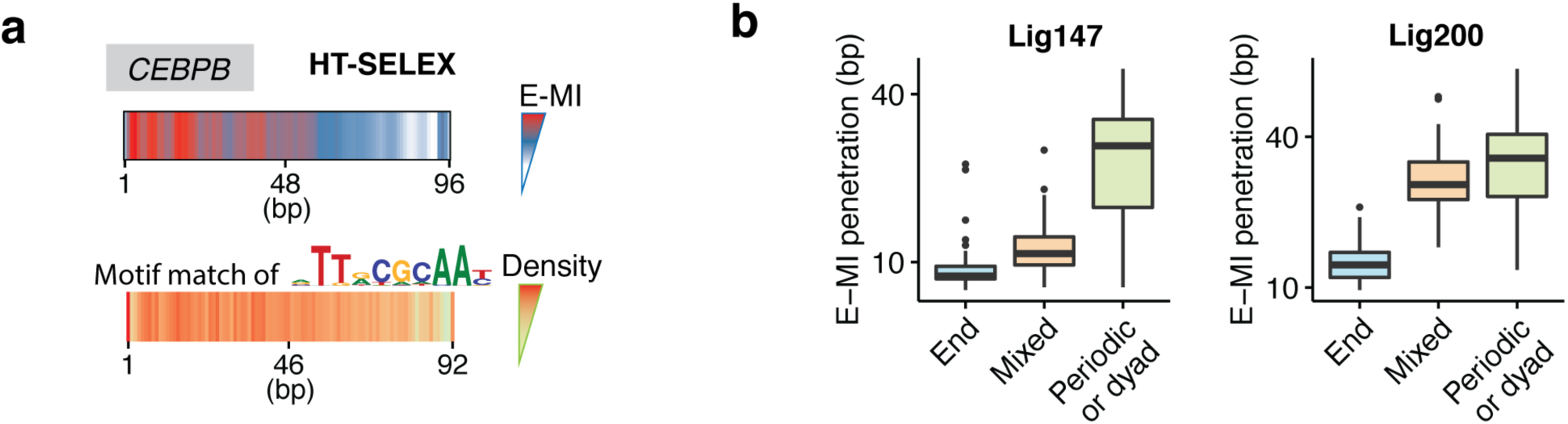
Control experiment for the end-binders and E-MI penetration according to binder classes. **a**, E-MI diagonal and motif matching results for the bZIP factor CEBPB in HT-SELEX. Without nucleosome, its signal distributes relatively homogeneously across the ligand. **b**, Penetration of E-MI signal for each binder class of TFs on lig147 and lig200. The results with SELEX ligands of different lengths generally correspond with each other. The periodic/dyad binders show deeper E-MI penetration than the end binders and thus are more capable of binding nucleosomal DNA. The boxes indicate the middle quartiles, separated by median line. Whiskers indicate last values within 1.5 times the interquartile range for the box.

**Extended Data Figure 6.**
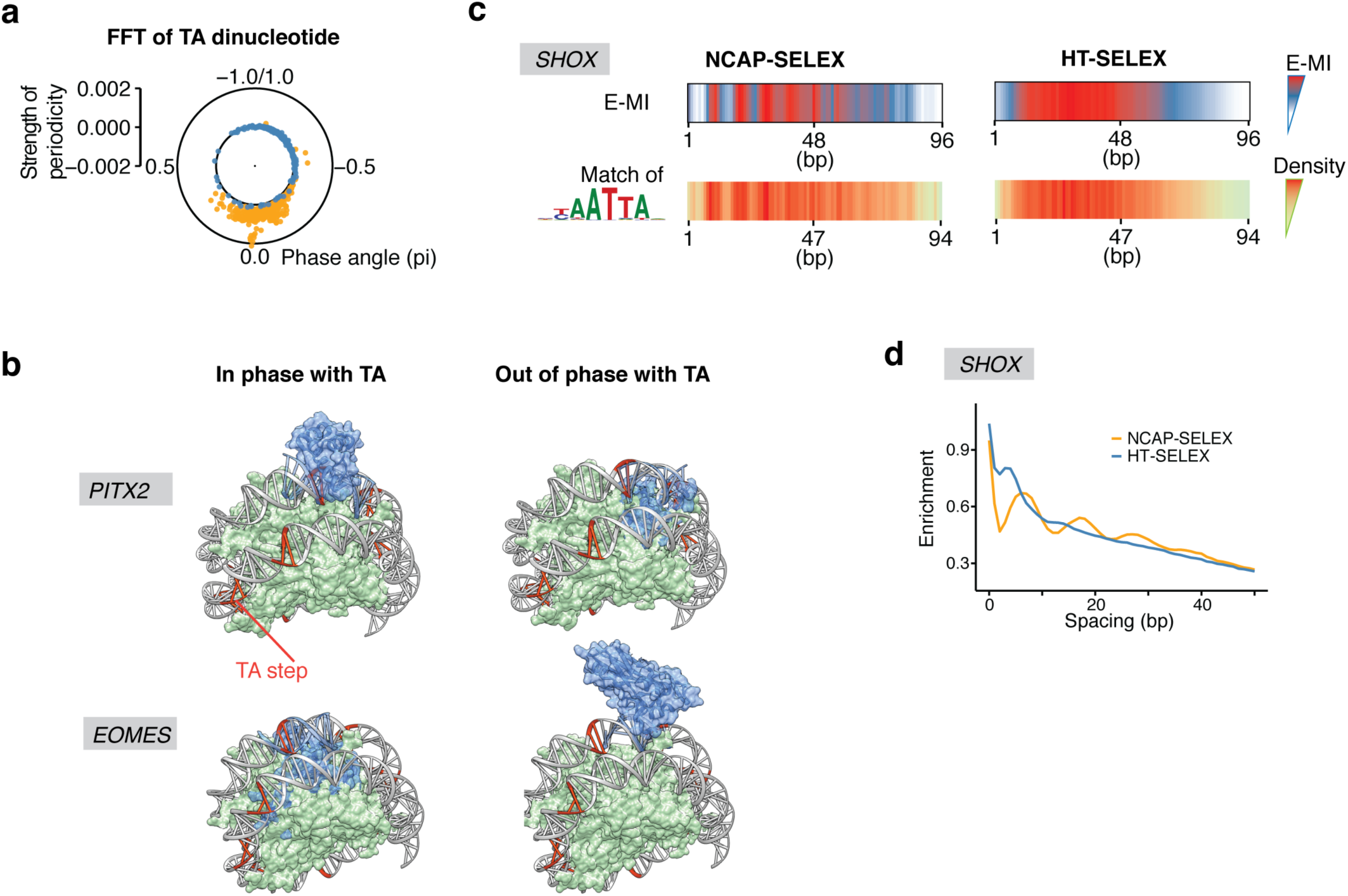
Analysis of the periodic binders. **a**, Strength and phase of the ~10 bp periodicity of TA dinucleotide in NCAP-SELEX and HT-SELEX libraries. For the library (lig147) enriched by a specific TF, the strength and phase information is derived from FFT of the TA counts at each position of the library. In the polar plot, each dot represents one TF’s library. The overall periodicity is stronger in the NCAP-SELEX libraries (yellow) than in the HT-SELEX libraries (blue), suggesting an enrichment of nucleosome signal. The TA phases in all TFs’ NCAP-SELEX libraries are similar, thus the rotational positioning of nucleosome on the SELEX ligand is similar for all TF’s libraries, **b**, Cartoon representations of the 3D structures of PITX3 (PDB 2lkx) and TBX5 (T_box, PDB 2×6 v) in complexes with nucleosomal DNA. The DNA ligand in the nucleosome structure (PDB 3ut9) contains phased TA steps (orange). Consistent with the SELEX result, PITX is more compatible with nucleosomal DNA when it binds in phase with TA, whereas T-box is more compatible when it binds out of phase with TA. **c**, E-MI diagonal and motif matching results for SHOX in NCAP-SELEX and HT-SELEX. **d**, The ~10 bp periodicity for the preferred spacing of SHOX dimers on nucleosomal DNA. In NCAP-SELEX libraries of many periodic binders (SHOX as an example), enrichment of the most abundant 3-mer tandem repeats oscillates as a function of the spacing between the repeats. The enrichment is evaluated by log2-ratio between the observed and expected occurrences.

**Extended Data Figure 7.**
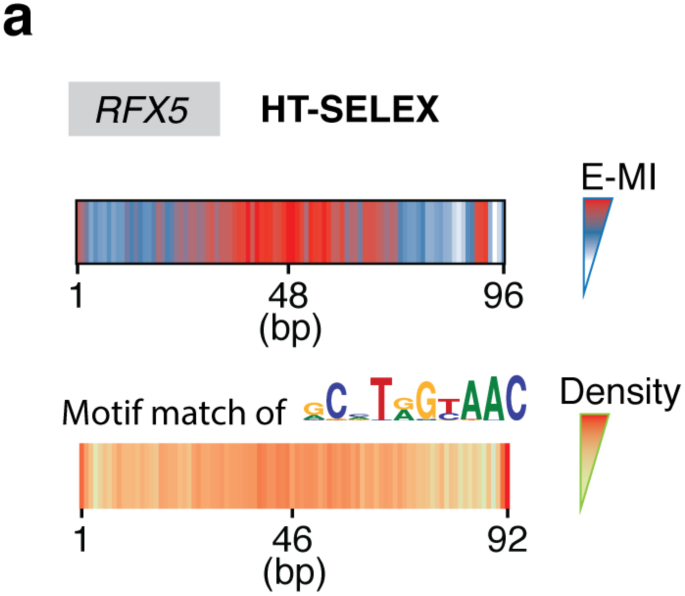
Analysis of the dyad binders. **a**, E-MI diagonal and motif matching results for RFX5 in HT-SELEX. The distribution of binding events is homogeneous in the absence of nucleosome.

**Extended Data Figure 8.**
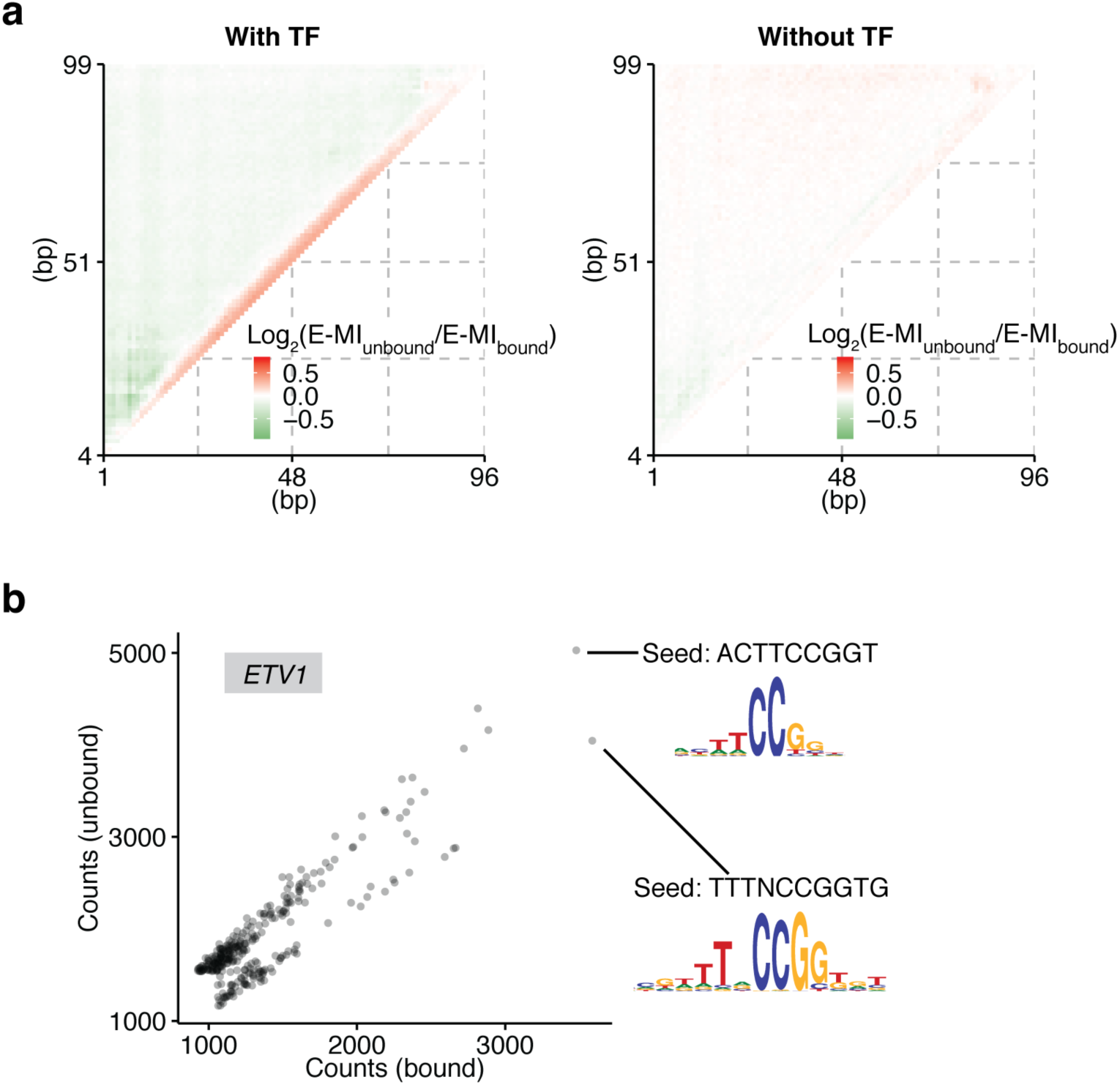
Effects of TF binding on the stability of nucleosome. **a**, E-MI difference between the bound and the unbound cycle **5** libraries. The bound and the unbound libraries were collected either in the presence (left) or in the absence (right) of TFs. The heatmaps visualize E-MI differences between the bound and unbound libraries for all position combinations of 3-mer pairs, and each pixel on the heatmap is a mean of all the examined TFs’ E-MI difference at this pixel. For individual TFs, value at each pixel is calculated as log2 (*E-MI_unbound,_/E-MI_bound_*). Testing nucleosome dissociation in the absence of TF was aimed to verify whether the TF motifs on lig147 by themselves can affect the nucleosome’s stability. **b**, The efficiency of nucleosome dissociation induced by ETV1 is dependent on its binding mode. The shorter mode is more efficient than the longer mode in displacing nucleosome, as it enriches more in the dissociated library (unbound).

